# A toolkit of highly selective and sensitive genetically encoded neuropeptide sensors

**DOI:** 10.1101/2022.03.26.485911

**Authors:** Huan Wang, Tongrui Qian, Yulin Zhao, Yizhou Zhuo, Chunling Wu, Takuya Osakada, Peng Chen, Huixia Ren, Yuqi Yan, Lan Geng, Shengwei Fu, Long Mei, Guochuan Li, Ling Wu, Yiwen Jiang, Weiran Qian, Wanling Peng, Min Xu, Ji Hu, Liangyi Chen, Chao Tang, Dayu Lin, Jiang-Ning Zhou, Yulong Li

## Abstract

Neuropeptides are key signaling molecules in the endocrine and nervous systems that regulate many critical physiological processes, including energy balance, sleep and circadian rhythms, stress, and social behaviors. Understanding the functions of neuropeptides *in vivo* requires the ability to monitor their dynamics with high specificity, sensitivity, and spatiotemporal resolution; however, this has been hindered by the lack of direct, sensitive and non-invasive tools. Here, we developed a series of GRAB (<u>G</u> protein-coupled <u>r</u>eceptor <u>a</u>ctivation‒<u>b</u>ased) sensors for detecting somatostatin (SST), cholecystokinin (CCK), corticotropin-releasing factor (CRF), neuropeptide Y (NPY), neurotensin (NTS), and vasoactive intestinal peptide (VIP). These fluorescent sensors utilize the corresponding GPCRs as the neuropeptide-sensing module with the insertion of a circular-permutated GFP as the optical reporter. This design detects the binding of specific neuropeptides at nanomolar concentration with a robust increase in fluorescence. We used these GRAB neuropeptide sensors to measure the spatiotemporal dynamics of endogenous SST release in isolated pancreatic islets and to detect the release of both CCK and CRF in acute brain slices. Moreover, we detect endogenous CRF release induced by stressful experiences *in vivo* using fiber photometry and 2-photon imaging in mice. Together, these new sensors establish a robust toolkit for studying the release, function, and regulation of neuropeptides under both physiological and pathophysiological conditions.

## INTRODUCTION

Neuropeptides were first identified nearly seven decades ago as hormone regulators in the endocrine system and have since been recognized as highly effective signaling molecules in both central and peripheral tissues (Du Vigneaud, 1954; Schally et al., 1973; Spiess et al., 1981; Wied and Kloet, 1987). In the brain, neuropeptides regulate many types of physiological functions, such as digestion, metabolism, sleep and circadian rhythm, reproduction, and higher cognitive processes (Alhadeff et al., 2018; Brazeau et al., 1973; de Lecea et al., 1998; Sakurai et al., 1998; Vale et al., 1981). For example, corticotropin-releasing factor (CRF) release orchestrates the responses to stress, and CRF hyperactivity increases arousal, alters locomotion, and decreases sexual receptivity and food consumption (Binder and Nemeroff, 2010; Dedic et al., 2018; Henckens et al., 2016). Neuropeptide Y (NPY) is enriched in the arcuate nucleus and is essential for stimulating food intake (Zhang et al., 2019); it also acts as an endogenous anticonvulsant in both rodents and humans (Baraban et al., 1997; Cattaneo et al., 2020; Colmers and El Bahh, 2003). Interestingly, neuropeptides, such as ghrelin, NPY, and neurotensin (NTS), were shown to exert neuroprotective effects in Parkinson’s disease and Alzheimer’s disease (Li et al., 2019; Zheng et al., 2021). Thus, neuropeptide signaling—which is mediated primarily by G protein-coupled receptors (GPCRs)—provides a key site for drug targeting for a wide range of diseases and conditions such as insomnia, pain, obesity, and diabetes (Davenport et al., 2020; Hauser et al., 2017; Hokfelt et al., 2003; Uslaner et al., 2013).

The ability to measure the spatial and temporal dynamics of neuropeptides *in vivo* is essential for understanding their functions and the mechanisms that regulate these key signaling molecules. However, current methods for detecting peptides in the brain either lack the necessary spatiotemporal resolution or are not suitable for *in vivo* application. Microdialysis of extracellular fluids combined with either an antibody-based radioimmunoassay or HPLC-MS requires large amounts of samples with long collection time and poor spatial precision (Al-Hasani et al., 2018; Andren and Caprioli, 1999; Ludwig et al., 2002; Merlo Pich et al., 1995). Reporter gene-based methods such as the Tango assay require at least several hours of reporter gene expression (Barnea et al., 2008; Lee et al., 2017; Mignocchi et al., 2020; Valtcheva et al., 2021). Fast-scan cyclic voltammetry offers higher temporal resolution but has so far only been used successfully in detecting one neuropeptide methionine enkephalin (Calhoun et al., 2019). Peptides tagged with large fluorescent proteins or reporters such as EGFP, pHluorin, and GCaMP provide a relatively fast readout of peptide release and describe knockout phenotypes related to peptide release *in vitro*, but this approach is limited to modified peptides, not the endogenous peptides, and is hardly possible to apply *in vivo* (Burke et al., 1997; de Wit et al., 2009; Ding et al., 2019; Lang et al., 1997; van den Pol, 2012; Xia et al., 2009). Finally cell-based neurotransmitter fluorescent engineered reporters (CNiFERs) have been used to detect neuropeptides; however, the need to implant exogenous cells in specific brain regions limits the utility of this approach (Jones et al., 2018; Lacin et al., 2016; Xiong et al., 2021). Thus, the precise spatiotemporal dynamics and release patterns of endogenous peptides remain poorly understood.

Genetically encoded fluorescent indicators have proven suitable for measuring the dynamics of signaling molecules with high spatiotemporal resolution *in vivo*. For example, bacterial periplasmic-binding protein (PBP)‒based sensors have been developed to detect neurotransmitters such as glutamate, acetylcholine, and serotonin (Borden et al., 2020; Marvin et al., 2013; Unger et al., 2020). However, corresponding PBPs for peptides and proteins are unlikely to exist thus generating peptide-sensing PBPs with high affinity and selectivity will require significant bioengineering and screening. Notably, most neuropeptide receptors are GPCRs, and peptide/protein GPCR ligands comprise 70% of all non-olfactory GPCR ligands in the human body (Figure 1A) (Foster et al., 2019; Isberg et al., 2016). Peptide GPCRs, thus, can be an ideal scaffold for binding and detecting specific peptide ligands, providing a valuable opportunity for generating genetically encoded sensors with high sensitivity and selectivity. Previously, our group and others developed and characterized several GPCR activation-based (GRAB) intensiometric biosensors, using GPCRs as the ligand-sensing unit and circular-permutated green fluorescent protein (cpGFP) as the reporter module (Feng et al., 2019; Jing et al., 2020; Jing et al., 2018; Patriarchi et al., 2018; Sun et al., 2018; Sun et al., 2020). The strategy for developing these GRAB sensors includes screening for the optimal cpGFP placement site within the receptor’s third intracellular loop 3 (ICL3); however, given the large number of peptide and protein GPCRs (with 131 expressed in humans) and the high variability of ICL3 among GPCRs (ranging from 2 to 211 amino acids), developing and optimizing a GRAB sensor for each GPCR would be highly labor-intensive (Otaki and Firestein, 2001; Unal and Karnik, 2012). Importantly, despite this structural variation in the ICL3, peptide GPCRs undergo a common structural change upon activation, with an outward movement of transmembrane 6 (TM6) observed in both class A and class B1 peptide GPCRs (Hollenstein et al., 2013; Ma et al., 2020; White et al., 2012) (Figure 1B). Thus, peptide GPCRs generated using the entire cpGFP-containing ICL3 in previously optimized GRAB sensors may retain the ability to couple the activation-induced conformational change with an increase in fluorescence, thereby accelerating the development of a wide variety of GRAB peptide sensors.

**Figure 1.**
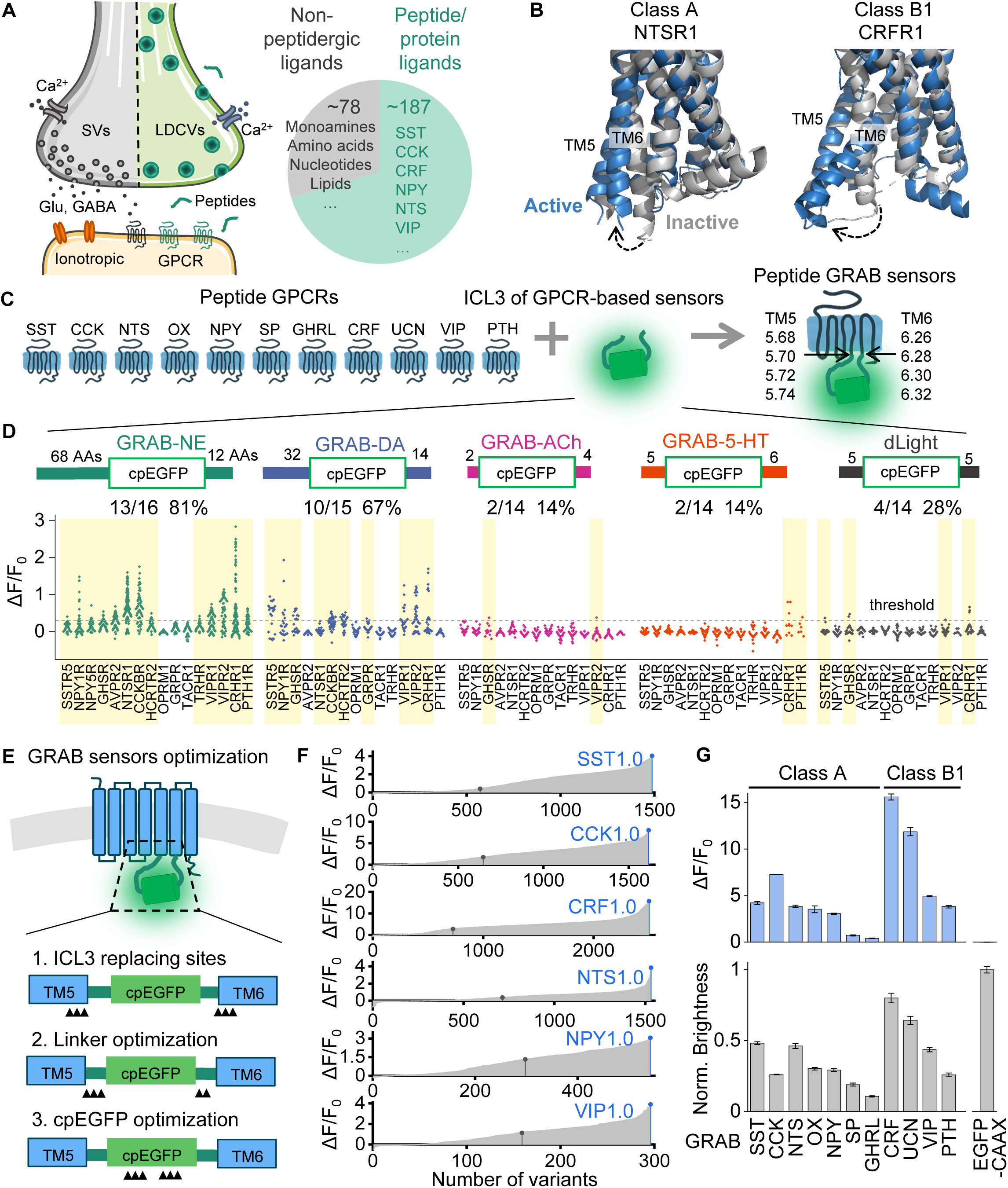
A general method for engineering fluorescent indicators for neuropeptides. (A) Left: illustration of peptide-containing large dense-core vesicles (LDCVs), neurotransmitter-containing synaptic vesicles (SVs), and their receptors in a synapse. Right: proportion and numbers of peptide/protein GPCR ligands and non-peptidergic GPCRs in humans, with corresponding examples. (B) Superposition of active (blue) and inactive (gray) structures of class A NTSR1 (PDB: 4BWB and 4GRV) and class B1 CRFR1 (PDB: 4K5Y and 6PB0). The dashed arrows indicate the movement of the sixth transmembrane domain (TM6). (C) Schematic diagram depicting the ICL3 transplantation strategy for developing GRAB sensors. (D) Fluorescence responses (ΔF/F_0_) of peptide GPCR chimeras with ICL3 transplanted from the indicated sensors. The amino acid numbers flanking cpGFP are labeled. The number and percentage of GPCRs with a maximum response exceeding 0.3 (dashed horizontal line) are shown, and these sensors are shaded in yellow. Each data point represents the average of 100-300 cells measured in one well. (E) Schematic diagram showing the steps for GRAB sensor optimization, including the ICL3 replacement site, linker optimization, and cpGFP optimization. Black triangles indicate the optimization sites. (F) Optimization of the SST, CCK, CRF, NTS, NPY, and VIP sensors. In each plot, the black dot indicates the initial version after ICL3 transplantation. After optimization, candidates with the highest ΔF/F0 were selected as the first-generation (1.0) sensors (blue dots). (G) Summary of the peak fluorescence response (top) and maximum brightness (bottom) of the indicated peptide sensors developed by transplanting ICL3 into the indicated class A and class B1 GPCRs (n = 4 wells containing 100-300 cells per well).

Here, we used this strategy to develop a series of GRAB sensors for detecting neuropeptides with ultra-high selectivity and nanomolar affinity. We then used these new peptide sensors to measure endogenous somatostatin (SST) release in isolated pancreatic islets, as well as cholecystokinin (CCK) and CRF release in acute brain slices with high spatiotemporal resolution. We also used these sensors to measure *in vivo* CRF levels using fiber photometry and 2-photon imaging during various behaviors. This new series of peptide sensors expands our repertoire of GRAB sensors, thus paving the way to addressing critical biological questions with respect to neuropeptide release and their roles under both physiological and pathophysiological conditions.

## RESULTS

### Developing a generalized method for engineering fluorescent sensors to detect neuropeptides

To develop a scalable method for generating a series of genetically encoded peptide sensors, we replaced the ICL3 domains in various peptide GPCRs with the ICL3 in several existing sensors, including GRAB_NE1m_, GRAB_DA2m_, GRAB_5- HT1.0_, GRAB_ACh3.0_, and dLight1.3b (Feng et al., 2019; Jing et al., 2020; Patriarchi et al., 2018; Sun et al., 2020; Wan et al., 2021). These sensor-derived ICL3s vary in length with respect to the number of amino acids that flank the cpGFP module (Figure S1A); thus, GRAB peptide sensors were generated by replacing the ICL3 in the GPCR with the linker sequences and cpGFP derived from the inner membrane regions of TM5 and TM6, located at sites around 5.70 and 6.28, respectively (Figure 1C). Each newly generated candidate peptide sensor was then expressed in HEK293T cells together with a plasma membrane-targeted mCherry (as a marker of surface expression). Each candidate’s performance was measured with respect to trafficking to the plasma membrane and the change in the sensor’s fluorescence in response to the appropriate ligand (Figures 1D and S1B-C). Candidates with a trafficking index over 80% (measured as the Pearsons correlation coefficient between the expression of a candidate and mCherry) and a fluorescence increase over 30% upon ligand application were considered as responsive peptide sensors.

We found peptide sensors containing the ICL3s derived from GRAB_NE1m_ and GRAB_DA2m_—both of which contain relatively long linkers (Figure S1A)—had a higher trafficking index and a larger response compared to sensors with ICL3s derived from GRAB_5-HT1.0_, GRAB_ACh3.0_, and dLight1.3b (Figure 1D). For further development, we chose the more generally applicable ICL3 in GRAB_NE1m_ and optimized the peptide sensor prototypes in four steps, including modifying the ICL3 donor sites, modifying the linker sequences, and modifying critical residues both in the cpGFP module and in the GPCR. The first three steps are depicted schematically in Figure 1E, and the whole optimization processes are shown for six peptide GRAB sensors in Figure 1F, in which the optimal version of the CRF sensor yielded a >10-fold increase in fluorescence upon CRF binding compared to the original candidate (Figures 1E-F and S1C). Utilizing the same strategy for both class A and class B1 peptide GPCRs—including the SSTR5, CCKBR, NTSR1, HCRTR2/OX2R, NPY1R, TACR1, GHS-R, CRF1R, CRF2R, VIPR2, and PTH1R receptors—we then developed and optimized a series of GRAB peptide sensors for detecting somatostatin (SST), cholecystokinin (CCK), neurotensin (NTS), orexin/hypocretin (OX), neuropeptide Y (NPY), substance P (SP), ghrelin (GHRL), corticotropin-releasing factor (CRF), urocortin (UCN), vasoactive intestinal peptide (VIP), and parathyroid hormone-related peptide (PTH) (Figures 1G and S1E).

### Characterization of GRAB peptide sensors in cultured cells

Next, we characterized the properties of the SST1.0, CCK1.0, CRF1.0, NPY1.0, NTS1.0, and VIP1.0 sensors. When expressed in HEK293T cells, all six GRAB sensors localized primarily to the plasma membrane and produced a robust change in fluorescence (ranging from a 2.5- to 12-fold increase in fluorescence) in response to their respective ligand (Figures 2A-B and S2C-D), and each response was blocked by the corresponding GPCR antagonist (Figures 2B and S2D). These sensors also retained the ligand selectivity of their respective GPCR scaffolds and had high sensitivity, with apparent half-maximum effective concentrations (EC_50_) of approximately 10-100 nM (Figures 2C, S1G, S2B, and S2E). As an example, the CRF1.0 sensor was based on the CRF1R receptor, which has a higher affinity for CRF than for urocortin (UCN) (Dautzenberg et al., 2004). As expected, the CRF1.0 sensor’s EC_50_ for CRF was 33 nM, compared to 68 nM for urocortin 1 (UCN1), while the peptides UCN2 and UCN3 had no effect on the CRF1.0 sensor (Figures 2C3 and S1G). We then tested the ligand specificity of these sensors and found that none of the sensors responded to glutamate (Glu), γ-aminobutyric acid (GABA), dopamine (DA), or any other neuropeptides tested (Figures 2D and S2A).

**Figure 2.**
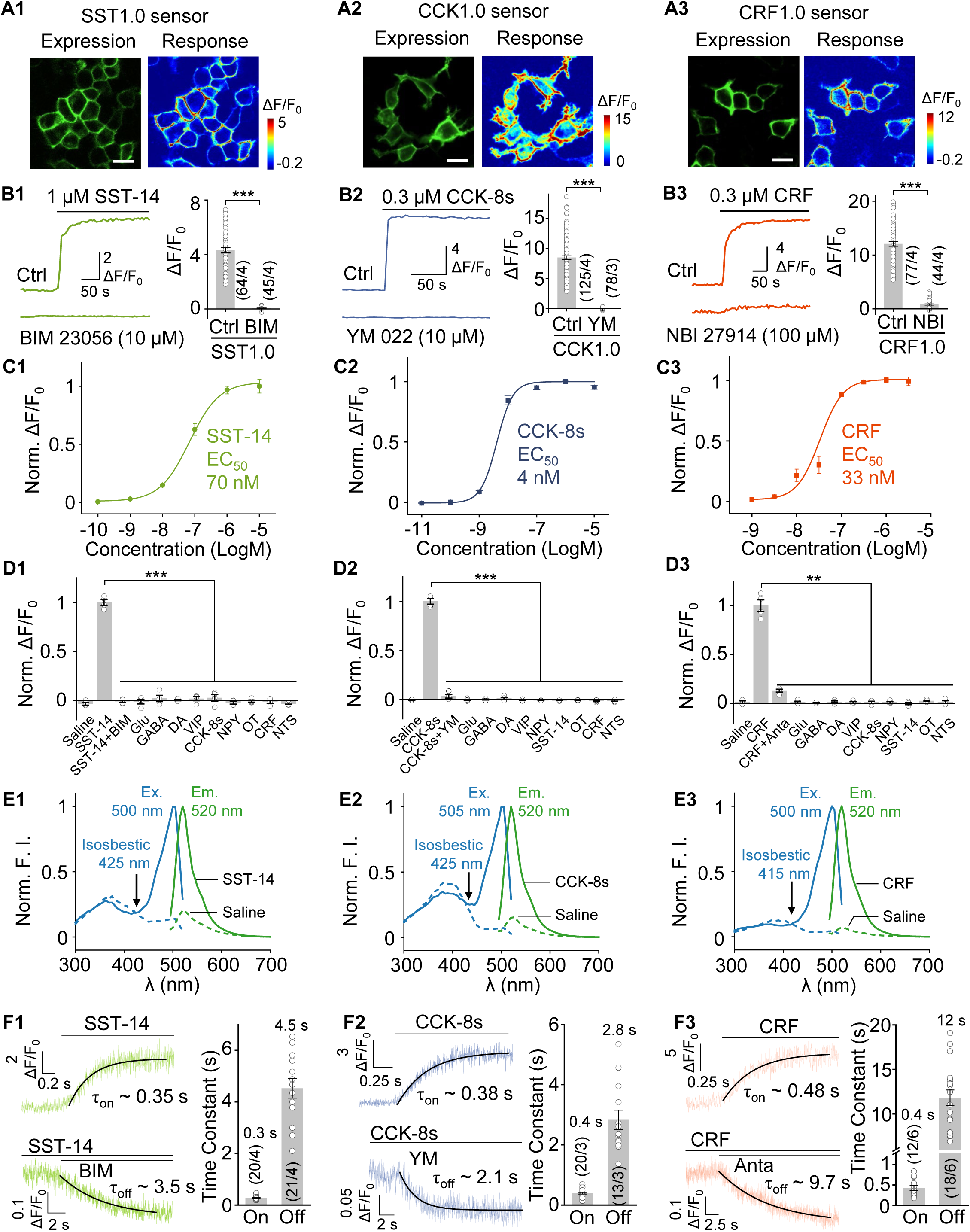
Characterization of the SST, CCK, and CRF sensors in HEK293T cells. (A) Representative images of HEK293T cells expressing SST1.0 (A1), CCK1.0 (A2), or CRF1.0 (A3), and the response to the application of SST-14 (1 μM), CCK-8s (300 nM), and CRF (300 nM), respectively. Scale bars, 20 μm. (B) Example fluorescence traces (left) and summary data (right) of HEK293T cells expressing SST1.0 (B1), CCK1.0 (B2), or CRF1.0 (B3); where indicated, the cells were pre-incubated with saline (Ctrl) or the antagonist BIM 23056 (10 μM), YM 022 (10 μM), or NBI 27914 (100 μM); n = 44-125 cells from 3-4 coverslips. (C) Normalized dose-response curves of HEK293T cells expressing SST1.0 (C1), CCK1.0 (C2), or CRF1.0 (C3) in response to the respective ligand (n = 3 wells containing 100-300 cells per well). (D) Summary of normalized ΔF/F_0_ in HEK293T cells expressing SST1.0 (D1), CCK1.0 (D2), or CRF1.0 (D3) in response to the indicated compounds applied; SST-14, CCK-8s, CRF, vasoactive intestinal peptide (VIP), neuropeptide Y (NPY), oxytocin (OT), and neurotensin (NTS) were applied at 1 μM, while Antalarmin (Anta), BIM 23056 (BIM), YM 022 (YM), glutamate (Glu), γ-aminobutyric acid (GABA), and dopamine (DA) were applied at 10 μM (n = 4 wells containing 100-300 cells per well). (E) Excitation (Ex) and emission (Em) spectra of SST1.0 (E1), CCK1.0 (E2), and CRF1.0 (E3) expressed in HEK293T cells, measured in the presence (solid curves) and absence (dashed curves) of SST-14 (10 μM), CCK-8s (1 μM), and CRF (1 μM), respectively. (F) Representative traces (left) and summary of τ_on_ and τ_off_ (right) of the SST1.0 (F1), CCK1.0 (F2), and CRF1.0 (F3) response. The indicated ligands and antagonists were locally puffed onto sensor-expressing cells, and high-speed line scanning was used to measure the fluorescence response. Where indicated, SST-14 (1 mM), CCK-8s (1 μM) and CRF (100 μM) for measuring signal increase; BIM 23056 (100 μM), YM 022 (30 μM), Antalarmin (100 μM), SST-14 (1 μM), CCK-8s (30 nM), and CRF (100 nM) were applied (n = 12-21 cells from 3-6 cultures).

We also measured the 1-photon spectra of these six peptide sensors and found a common excitation peak at ∼500 nm and a common emission peak at ∼520 nm, with an isosbestic point of the excitation wavelength at ∼420 nm (Figures 2E and S2F). The 2-photon excitation cross-section of the SST, CCK, and CRF sensors showed excitation peaks at 920-930 nm in the presence of the respective ligands (Figure S1F). The kinetics of the peptide sensors’ responses were also measured by locally applying the corresponding peptide ligands and antagonists and then recording the change in fluorescence using line-scan confocal microscopy. The resulting time constants of the rise in the signal (τ_on_) ranged from approximately 0.3 s to 0.9 s, and the time constants of the signal decay (τ_off_) ranged from approximately 3 s to 12 s (Figures 2F and S2G).

Next, we measured the properties of our GRAB peptide sensors expressed in cultured rat cortical neurons. Consistent with their expression in HEK293T cells, the sensors localized to the neuronal membrane both at the cell body and in extended ramified neurites, and responded robustly to ligand application (Figures 3A and S3A). Moreover, when expressed in cultured neurons, the peptide sensors’ responses and apparent EC_50_ values were similar to those measured in HEK293T cells, and the responses were again blocked by the respective GPCR antagonists (Figures 3B-C and S3B-C). Finally, for most of the peptide sensors, the ligand-induced change in fluorescence was stable for up to 120 min in neurons exposed to saturated ligand concentration (Figures 3D and S3D), indicating that the peptide sensors have minimal internalization, even during chronic activation.

**Figure 3.**
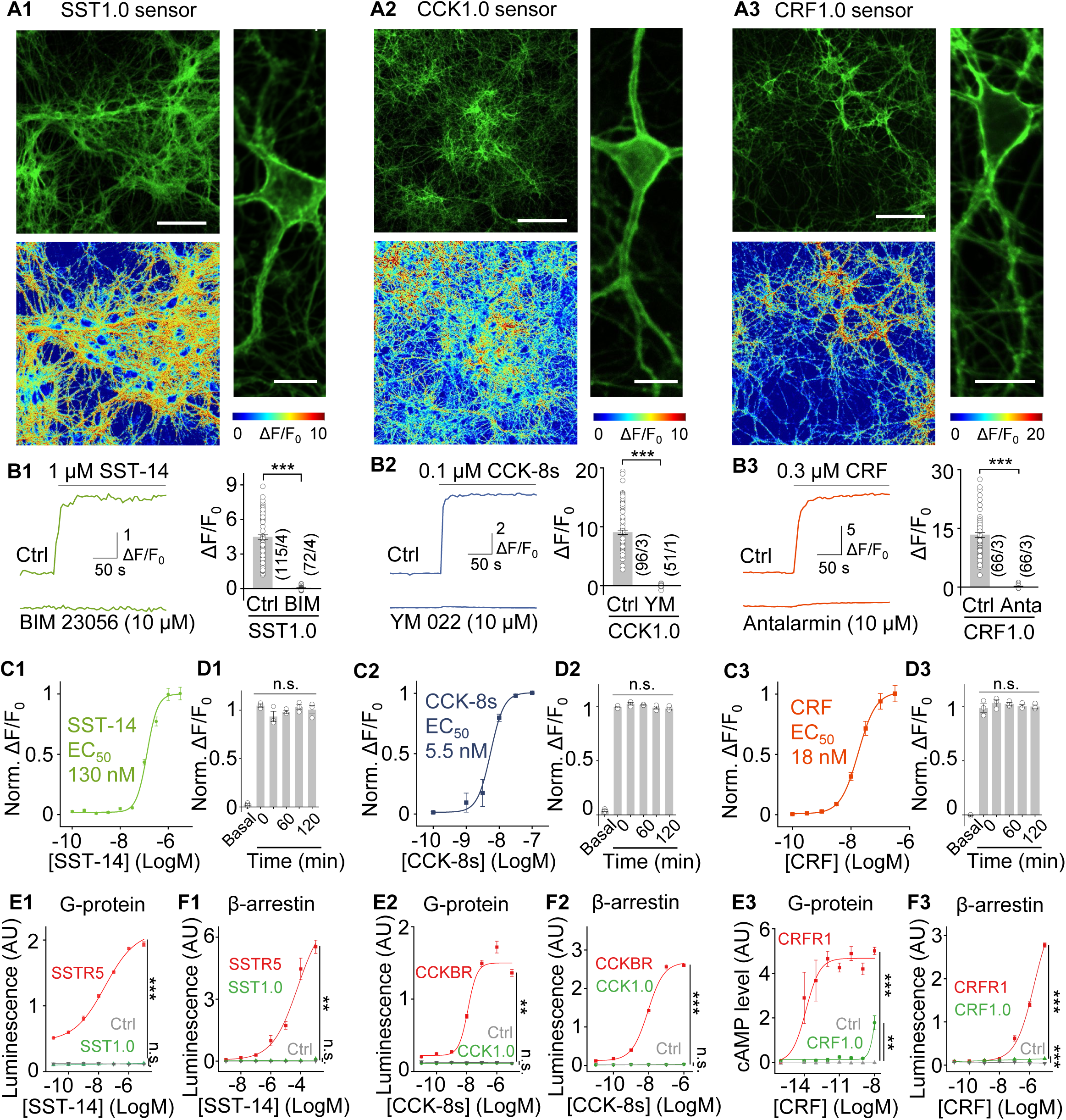
Characterization of the SST, CCK, and CRF sensors in cultured neurons and cell lines. (A) Representative images of primary cultured rat cortical neurons expressing SST1.0 (A1), CCK1.0 (A2), or CRF1.0 (A3), showing sensor expression (top left), pseudocolor responses (bottom left), and cell membrane localization (right). Scale bars represent 100 μm (left) and 20 μm (right). (B) Example fluorescence traces (left) and summary data (right) of neurons expressing SST1.0 (B1), CCK1.0 (B2), or CRF1.0 (B3); where indicated, peptides and agonists were applied at 10 μM (n = 51-115 regions of interest (ROIs) from 1-4 coverslips. (C) Normalized dose-response curves of neurons expressing SST1.0 (C1), CCK1.0 (C2), or CRF1.0 (C3) in response to the indicated ligands; n = 3 cultures each with 20-40 ROIs. (D) Summary of the fluorescence change measured in neurons expressing SST1.0 (D1), CCK1.0 (D2), or CRF1.0 (D3) in response to a 2-hour continuous application of 1 μM SST-14, 100 nM CCK-8s, or 300 nM CRF, respectively; n = 4 cultures each with 20-40 ROIs. (E and F) G protein and β-arrestin coupling were measured using the split-luciferase complementation assay (E1 and E2), a cAMP reporter (E3), and the Tango assay (F1-F3) in cells expressing either the wild-type peptide receptor (red), sensor (green), or no receptor (Ctrl; gray) in the presence of the indicated concentrations of the ligand; n = 3 wells each.

We then tested whether our GRAB peptide sensors can couple to downstream signaling pathways by measuring G protein-mediated signaling and beta-arrestin recruitment. Although wild-type peptide receptors activated both signaling pathways, their corresponding GRAB sensors elicited significantly reduced or virtually no downstream signaling (Figure 3E-F), highlighting that overexpression of peptide sensors does not disrupt endogenous signaling.

Taken together, these results indicate that our SST, CCK, CRF, NPY, NTS, and VIP sensors are all highly sensitive, specific, and produce a robust real-time increase in fluorescence in response to their corresponding ligands, without activating downstream signaling pathways. We chose the SST, CCK, and CRF sensors for further study.

### The SST1.0 sensor can be used to detect the release of endogenous SST in cortical neurons

Neuropeptides are widely used as markers to categorize various types of neurons, with SST-expressing neurons representing subsets of GABAergic interneurons in the cerebral cortex (Smith et al., 2019; Tremblay et al., 2016; Wamsley and Fishell, 2017). Although used as a marker for neuronal subpopulations, whether SST is actually released from cortex neurons—and the spatiotemporal pattern of its potential release—has not been well investigated. Previous studies showed that applying trains of electrical field stimuli to cultured mouse hippocampal neurons can induce the fusion of peptide-containing dense-core vesicles (Arora et al., 2017; Persoon et al., 2018). To detect SST release from these neurons, we expressed the SST1.0 sensor in cultured primary rat cortical neurons. We found that applying increasing numbers of pulse trains elicited increasingly strong responses (Figure 4A-C). Application of 75 mM K^+^ to depolarize the neurons also induced a robust increase in SST1.0 fluorescence that was blocked by the SST receptor antagonist BIM 23056; moreover, no increased response was measured in neurons expressing the membrane-targeted EGFP-CAAX (Figure 4A-C). The rise and decay half-times of the SST1.0 signal induced by stimulation and K^+^ application are summarized in Figure 4D. Furthermore, the SST1.0 response was directly correlated with the corresponding increase in cytosolic Ca^2+^ levels measured using the fluorescent Ca^2+^ indicator Calbryte-590 (Figure 4E). These results suggest that the SST1.0 sensor can reliably report the release of endogenous SST in cultured cortical neurons.

**Figure 4.**
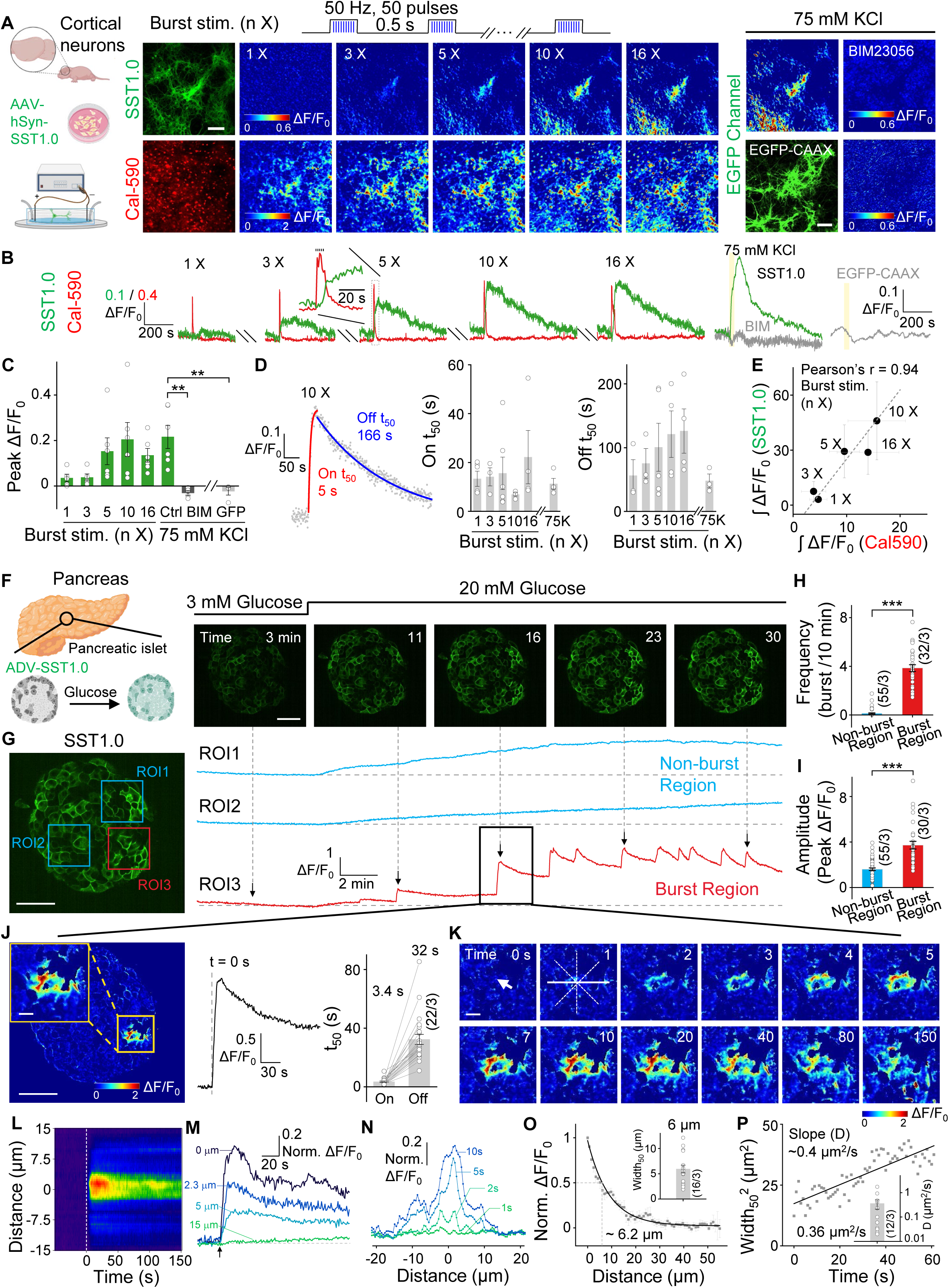
Imaging SST release in cultured neurons and pancreatic islets. (A) Left: schematic diagram depicting the experimental strategy. Middle: representative fluorescence images and pseudocolor images of rat cortical neurons both expressing SST1.0 (top row) and loaded with Calbryte-590 (bottom row); where indicated, burst electrical stimuli (50 pulses delivered at 50 Hz, 0.5 s interval between bursts) were applied. Right: representative fluorescence images of rat cortical neurons expressing SST1.0 (top row) or EGFP-CAAX (bottom row); 75 mM K^+^ was applied by perfusion in the absence (left column) or presence (right column) of the SST receptor antagonist BIM. Scale bars, 100 μm. (B) Example traces of the change in SST1.0 (green) and Calbryte-590 (red) in response to electric stimuli; SST1.0 (with or without antagonist BIM) and EGFP-CAAX fluorescence in response to 75 mM K^+^; yellow shadings indicate the 75 nM KCl perfusion time. (C) Summary of the peak change in fluorescence measured in neurons expressing SST1.0 or EGFP-CAAX in response to burst stimuli or 75 mM K^+^. (D) Representative time course (left) and summary of the rise and decay half-times (t_50_) of the fluorescence change (right) measured in SST1.0-expressing neurons in response to burst stimuli and 75 mM K^+^; n = 3-5 cultures containing 20-40 ROIs per culture. (E) Change in SST1.0 fluorescence plotted against the Calbryte-590 signal measured in response to the indicated number of burst stimuli. (F) Left: schematic diagram depicting the experimental strategy in which pancreatic islets were isolated, infected with adenoviruses expressing SST1.0, and treated with high (20 mM) glucose. Right: example fluorescence images of an SST1.0-expressing pancreatic islet before and after application of 20 mM glucose. Scale bar, 50 μm. (G) SST1.0 fluorescence was measured at the indicated ROIs in the same pancreatic islet shown in (F). Based on the response patterns (right panel), ROI1 and ROI2 are classified as non-burst regions (blue), while ROI3 is classified as a burst region (green). Scale bar, 50 μm. (H and I) Summary of the burst frequency (H) and peak response (I) measured of non-burst and burst regions; n = 30-55 ROIs from 3 islets. (J) Left: example pseudocolor image of maximum response taken at the indicated time from ROI3 in the pancreatic islet (G). Middle: the fluorescence response trace corresponding to the single burst at the indicated time. Right: summary of the rise and decay t_50_ values measured for 22 burst events in 3 islets. (K) Example time-lapse pseudocolor images of SST1.0 fluorescence measured in the burst region indicated by the yellow square in (J). The white arrow indicates the location from which the signal originates. Scale bar, 10 μm. (L-N) Representative spatial-temporal profile (L), temporal dynamics (M), and spatial dynamics (N) of the SST1.0 fluorescence response measured during a single burst. The arrow in (M) indicates time 0. The traces in (M) and (N) correspond to the indicated distances and times, respectively. (O) Normalized fluorescence responses measured at 10 s fitted with a single-exponential function, showing a signal width_50_ of approximately 6.2 μm. The inset shows the summary width_50_ data (n = 16 burst events from 3 islets). (P) Distribution of the (width_50_)^2^ over time, fitted to a linear function with a slope of 0.4 μm^2^/s. The inset summarizes the effective diffusion coefficient (D); note that the *y*-axis is a log scale (n = 12 burst events from 3 islets).

### The SST1.0 sensor can be used to detect glucose-stimulated SST release in isolated pancreatic islets

SST plays an essential role in feeding and energy expenditure by affecting central and peripheral tissues (Kumar and Singh, 2020). In pancreatic islets, the release of SST from delta (δ) cells (comprising ∼5% islets cells) is critical for regulating the activity of glucagon-releasing α cells and insulin-releasing β cells, thus playing a role in controlling blood glucose (Hauge-Evans et al., 2009; Rorsman and Huising, 2018); however, the spatiotemporal pattern of SST release in individual islets has not been investigated. To measure SST release in islets, we expressed SST1.0 under the control of a non-selective CMV promoter in mouse pancreatic islets cultures using adenovirus infection. We found that application of the peptide SST-14—but not CCK, which also exists in islets and stimulates pancreatic enzyme secretion (Rehfeld, 2017)—caused a robust increase in SST1.0 fluorescence, and this response was blocked by the SST receptor antagonist BIM 23056 but not the CCK receptor antagonist YM 022 (Figure S4A-C). We then examined whether SST1.0 can detect the release of endogenous SST in islets in response to high glucose stimulation (Hellman et al., 2012; Salehi et al., 2007). Application of 20 mM glucose caused a progressive increase in SST1.0 fluorescence (Figures 4F-G and S4D-F). Moreover, the increase in SST1.0 fluorescence had a distinct spatial pattern within the islet, with regions that could be classified as non-burst and burst regions (Figure 4G and Video S1). Analyzing these regions separately revealed that burst regions exhibited a phasic SST1.0 response in the presence of 20 mM glucose, with a higher burst rate and larger peak responses compared to non-burst regions (Figure 4H-I). Notably, during a single burst event, the SST1.0 signal first increased at a focal hotspot and then propagated over time to neighboring cells (Figure 4J-L). The response measured near the initial hotspot was more rapid and robust than the responses measured farther away from the hotspot (Figure 4M). Moreover, at time point 10 s, this propagation of the SST1.0 signal had an average half-width of ∼6 μm (Figure 4N-O), and this half-width increased over time, with an average diffusion coefficient of approximately 0.4 μm^2^/s (Figure 4P).

### Characterization of the CCK1.0 sensor expressed in acute brain slices

The peptide CCK is also expressed at high levels in the brain (Dockray, 1976; Muller et al., 1977; Vanderhaeghen et al., 1975), particularly in subpopulations of interneurons in the cortex and hippocampus (Beinfeld et al., 1981; Nunzi et al., 1985; Whissell et al., 2015). To examine whether our CCK1.0 sensor can be used to measure the release of endogenous CCK in the mouse brain, we injected adeno-associated virus (AAV) expressing CCK1.0 (or EGFP-CAAX as a negative control) into the hippocampus (Figure 5A). After 3 weeks of expression, we prepared coronal brain slices containing the hippocampal CA1 and applied electrical stimuli to activate the CA1 neurons (Figure 5B). We found that applying stimuli at 20 Hz induced increases in CCK1.0 fluorescence, and the magnitude of the peak response increased with the number of pulses. Moreover, the response was frequency-dependent, increasing as frequency increased from 2 Hz to 50 Hz. The response was blocked by treating the slices with the CCK receptor antagonist YM 022, and no response was detected in EGFP-CAAX‒expressing CA1 slices (Figure 5C-E). The rise and decay half-times of the CCK1.0 response also increased with pulse numbers and were 0.8-4.7 s and 2.0-5.0 s, respectively (Figure 5F). Propagation of the stimulation-induced CCK1.0 response was also observed (Figure 5G-J), with an apparent diffusion coefficient of 1.7 x 10^3^ μm^2^/s (Figure 5K-L).

**Figure 5.**
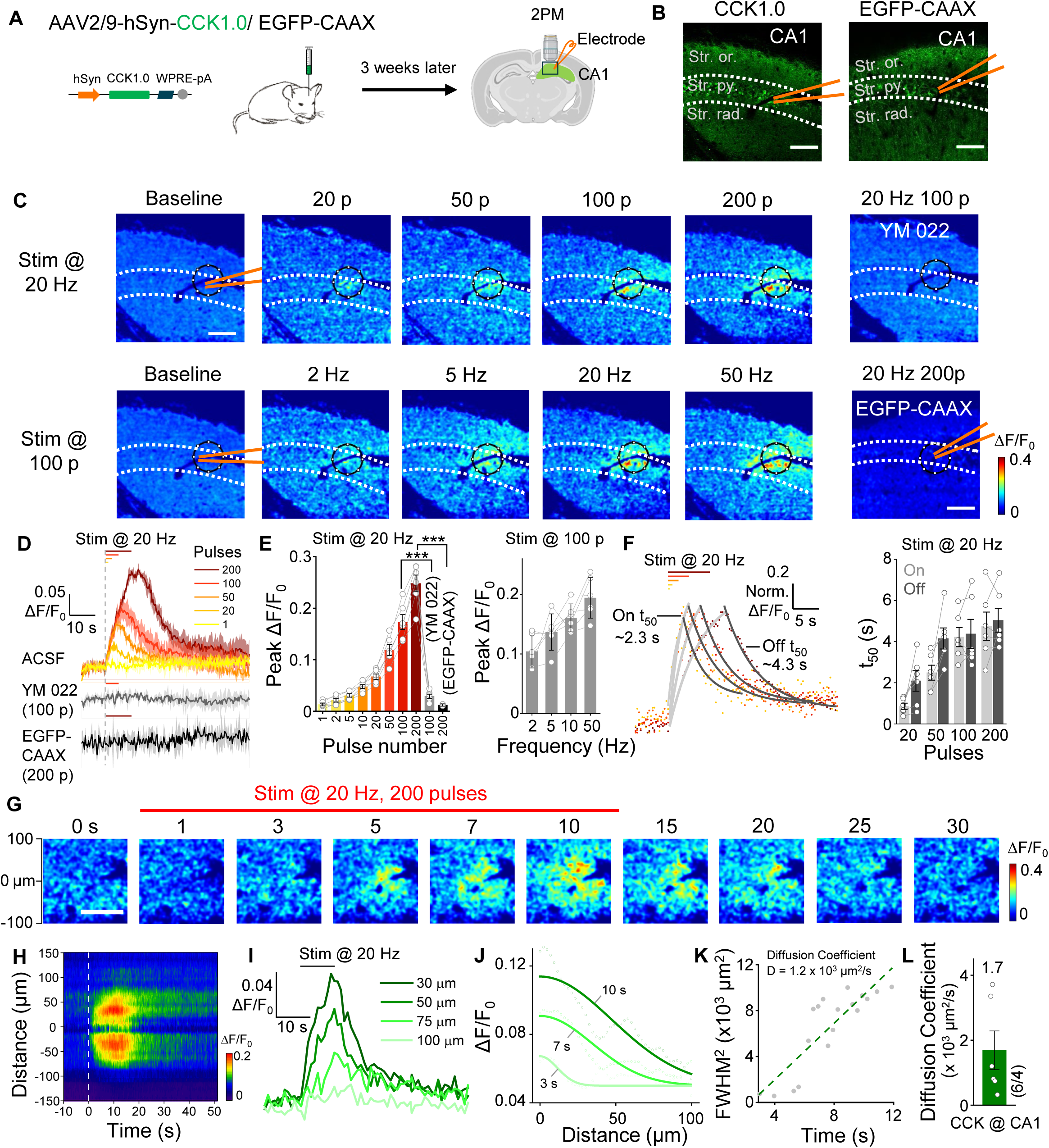
Validation of the CCK1.0 sensor in acute hippocampal slices. (A) Schematic illustration depicting the experimental design in which CCK1.0 or EGFP-CAAX was virally expressed in the mouse hippocampus; after 3 weeks, acute slices were prepared. (B) Representative fluorescence images showing CCK1.0 or EGFP-CAAX expression in the CA1 region; the location of the stratum pyramidale, stratum oriens, and stratum radiatum are indicated, and the approximate location of the stimulating electrode is shown. Scale bars, 100 μm. (C) Example pseudocolor images of CA1 slices expressing CCK1.measured at baseline and in response to 20, 50, 100, and 200 pulses delivered at 20 Hz (top row), or 100 pulses delivered at 2, 5, 20, and 50 Hz (bottom row). The images at the right show a slice treated with 10 μM YM 022 and stimulated with 100 pulses at 20 Hz (top) and a slice expressing EGFP-CAAX and stimulated with 200 pulses at 20 Hz (bottom). The black circles indicate the ROI used to analyze the responses. (D-E) Representative traces (D) and summary (E) of the change in CCK1.0 fluorescence in response to various numbers of electric stimuli delivered in ACSF, 100 pulses delivered in YM 022, and the change in EGFP-CAAX fluorescence in response to 200 pulses. Shown at the right is the peak change in CCK1.0 fluorescence in response to 100 pulses delivered at the indicated frequencies. n = 2-6 slices from 1-4 mice. (F) Representative fitted curves (left) and summary (right) of on and off t_50_ of the change in CCK1.0 fluorescence; n= 6 slices from 4 mice. (G) Example time-lapse pseudocolor images of CAI slices expressing CCK1.0; during the first 10 s, 200 pulses were applied at 20 Hz. Scale bar, 100 μm. (H-J) Representative spatial-temporal profile (H), temporal dynamics (I), and spatial dynamics (J) of the fluorescence change shown in (G). The profile in (H) shows the average response of three trials conducted in one slice. The traces in (I) and (J) correspond to the indicated distances and times, respectively, and the data in (J) were fitted with a Gaussian function. (K) Representative plot of FWHM^2^ (full width at half maximum, squared) against time using the data shown in (J); the diffusion coefficient (D) was measured as the slope of a line fitted to the data. (L) Summary of the diffusion coefficient (D) for CCK measured in the CA1 region; n = 6 slices from 4 mice.

Finally, we found that the CCK1.0 sensor could be expressed in the hippocampal CA1 region and responded to bath application of 75 mM K^+^ and 1 μM CCK-8s (Figure S4G-I). These results indicate that our CCK sensor can report the release of endogenous CCK in acute brain slices with relatively good temporal and spatial resolution.

### The CRF1.0 sensor can be used to measure CRF release in acute brain slices and *in vivo*

CRF is considered to be an anxiogenic neuropeptide, and CRF neurons in the central amygdala (CeA) play an important role in several conditions related to fear, anxiety, and alcohol addiction (de Guglielmo et al., 2019; Flandreau et al., 2012; Jo et al., 2020; Pomrenze et al., 2019; Regev et al., 2011; Sanford et al., 2017). To test whether the CRF1.0 sensor can be used to measure the release of endogenous CRF in the CeA, we expressed either the CRF1.0 sensor in the CeA and then recorded the response in acute brain slices using 2-photon fluorescence microscopy (Figure 6A-B). We found that electric stimuli delivered at 20 Hz induced a robust increase in CRF1.0 fluorescence, with larger responses induced by increased numbers of pulses, and that this response was significantly blocked by treating the slices with the CRF receptor antagonist AHCRF (alpha-helical CRF) (Figure 6C-D); in contrast, no response was measured in slices expressing EGFP-CAAX (Figure 6D). We also found that the rise and decay half-times increased with increasing pulse numbers, with on and off t_50_ values of approximately 0.6-1.8 s and 3.5-6.4 s, respectively (Figure 6E). Finally, we found that the CRF1.0 signal propagated during electrical stimulation (Figure 6F-I), with an average diffusion coefficient of 3.5×10^3^ μm^2^/s (Figure 6J-K). These results indicate that the CRF1.0 sensor can report the release of endogenous CRF in acute brain slices with relatively good temporal and spatial resolution.

**Figure 6.**
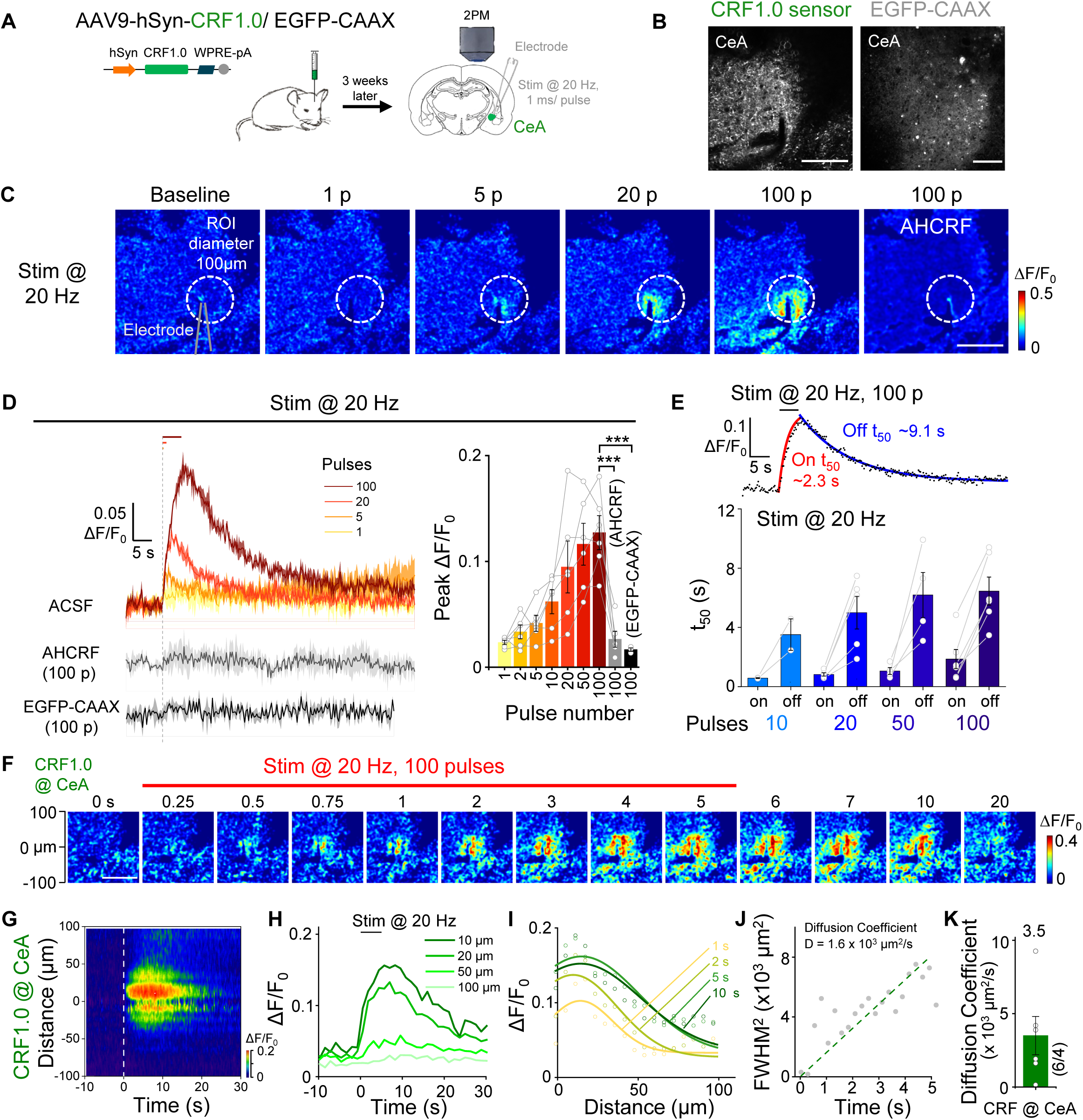
Detection of endogenous CRF release in acute brain slices using CRF1.0. (A) Schematic illustration depicting the experimental design in which CRF1.0 or EGFP-CAAX was expressed virally in the central amygdala (CeA); after 3 weeks, acute slices were prepared. (B) Representative 2-photon fluorescence images of acute slices, showing expression of CRF1.0 and EGFP-CAAX in the CeA. Scale bars, 100 μm. (C) Example pseudocolor images of acute slices expressing CRF1.0- at baseline and in response to 1, 5, 20, and 100 electric stimuli delivered at 20 Hz, and the response to 100 pulses measured in the presence of 100 nM alpha-helical CRF (AHCRF). The dashed white circles indicate the ROI used to calculate the response, and the approximate position of the stimulating electrode is indicated. (D) Representative traces (left) and summary (right) of the change in CRF1.0 fluorescence in response to electric stimuli delivered at 20 Hz in ACSF and 100 pulses delivered in the presence of AHCRF; also shown is the response measured in slices expressing EGFP-CAAX. n = 3-6 slices from 1-3 mice. (E) Representative fitted curves (left) and summary (right) of on and off t_50_ of the change in CRF1.0 fluorescence; n = 2-6 slices. (F) Example time-lapse pseudocolor images of CRF1.0 expressed in the CeA; during the first 5 s, 100 pulses were delivered at 20 Hz. Scale bar, 100 μm. (G-I) Representative spatial-temporal profile (G), temporal dynamics (H), and spatial dynamics (I) of the fluorescence change shown in (F). The profile in (G) shows the average response of three trials conducted in one slice. The traces in (H) and (I) correspond to the indicated distances and times, respectively, and the data in (I) were fitted with a Gaussian function. (J) FWHM^2^ plotted against time based on the data shown in (I); the diffusion coefficient (D) was measured as the slope of a line fitted to the data. (K) Summary of the diffusion coefficient (D) measured CRF in the CeA; n = 6 slices from 3 mice.

CRF neurons in the paraventricular nucleus of the hypothalamus (PVN) have been known to play an essential role in regulating the stress response via the endocrine axis (Vale et al., 1981). In addition, these neurons also respond rapidly to both aversive and appetitive stimuli (Daviu et al., 2020; Kim et al., 2019; Yuan et al., 2019). To investigate the specificity of our CRF1.0 sensor *in vivo*, we expressed CRF1.0 or a CRF-insensitive mutant (CRFmut) (see Figure S5A-D for expression in HEK293T cells) in the mouse PVN. We recorded the signal using fiber photometry while infusing CRF and/or AHCRF via an intracerebroventricular cannula (Figure 7A). We found that CRF1.0 fluorescence increased in a dose-dependent manner after CRF infusion (Figure 7B), and the increase was blocked by co-administration of AHCRF (Figure 7D); in contrast, CRFmut expressed in the PVN showed virtually no response to CRF, even at the highest concentration (Figure 7C).

**Figure 7.**
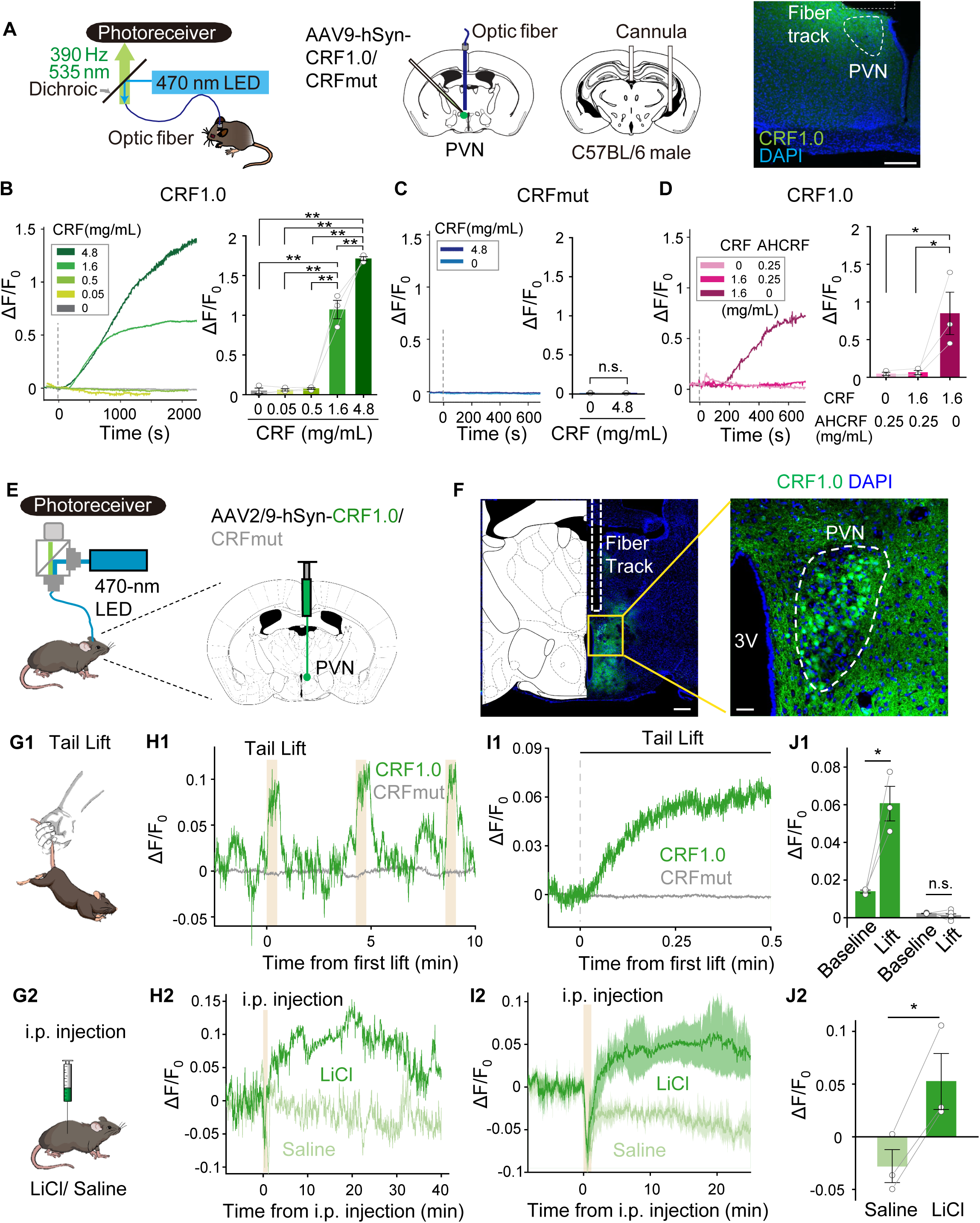
Using fiber photometry to measure endogenous CRF release *in vivo*. (A) Left: schematic diagrams depicting the strategy for virus injection and fiber and cannula implantation, and measurement of CFR1.0 or CRFmut in the paraventricular nucleus (PVN). Right: image showing the expression of CRF1.0 (green) in the PVN and the approximate location of the optic fiber above the PVN; the nuclei were counterstained with DAPI (blue). Scale bar, 200 μm. (B-D) Representative traces (left panels) and summary of the response (right panels) measured in mice expressing CRF1.0 (B and D) or CRFmut (C); the indicated concentrations of CRF and alpha-helical CRF 9-41 (AHCRF) were infused via the cannula. (E) Schematic diagram depicting the strategy for virus injection and fiber photometry recording. (F) Image showing the expression of CRF1.0 (green) and the approximate location of the imaging fiber; the nuclei were counterstained with DAPI (blue). Scale bars, 300 μm (left) and 40 μm (right). (G-J) Illustration (G), representative traces (H), average traces per stimulus-response (I), and summary data (J) of the change in CRF1.0 and CRFmut fluorescence measured before and during a 30-s tail lift (G1-J1) and before and after an i.p. injection of LiCl or saline (G2-J2); n = 3-6 animals.

Next, we measured the dynamics of CRF release in the PVN during stressful experiences in mice expressing CRF1.0 (Figures 7E-F). We found that suspending the mouse by the tail for 30 s induced a robust time-locked increase in CRF1.0 fluorescence, while mice expressing CRFmut or EGFP-CAAX in the PVN showed no visible response (Figure 7G1-J1 and S5E-H). Similarly, an i.p. injection of LiCl, an abdominal malaise-inducing stimulus, but not saline, elicited a long-lasting increase in CRF1.0 fluorescence, while no response was measured in mice expressing CRFmut or EGFP-CAAX (Figures 7G2-J2 and S5E-H, the rise halftimes are shown in Figure S5I). Taken together, these results indicate that various stress-inducing stimuli trigger the release of CRF in the PVN, and this release can be measured using CRF1.0 *in vivo* in real-time.

CRF is expressed abundantly in neocortical interneurons, and CRF receptors are present in pyramidal cells (Deussing and Chen, 2018; Gallopin et al., 2006). In the frontal cortex, CRF mediates stress-induced executive dysfunction (Chen et al., 2020; Uribe-Marino et al., 2016; Zięba et al., 2008). We, therefore, investigated the role of CRF in the mouse cortex during various behavioral paradigms. We injected virus expressing CRF1.0 into the motor cortex and prefrontal cortex (PFC) and then performed 2-photon imaging of CRF1.0-expressing layer 2/3 neurons in head-fixed mice (Figure 8A). We observed a transient increase in CRF1.0 fluorescence in both the motor cortex and PFC in response to tail shocks; in contrast, no response was detected in mice expressing CRFmut or EGFP-CAAX (Figure 8B1-D1, G and Figure S6A-E; the kinetics and time constants are shown in Figure 8E-F).

**Figure 8.**
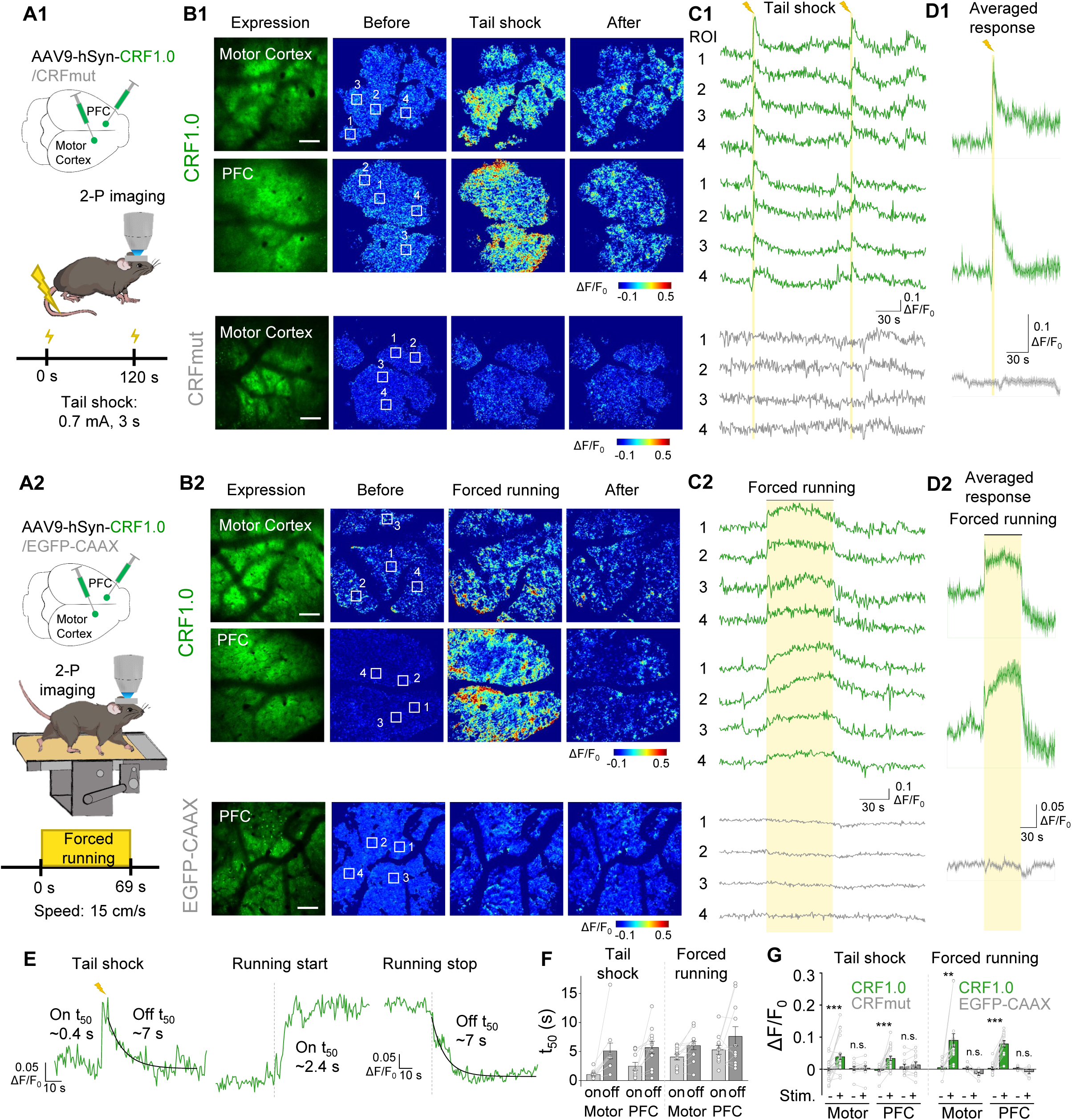
Spatially resolved measurements of CRF *in vivo* using two-photon imaging. (A-D) Schematic diagrams (A), representative expression and pseudocolor response images (B), representative traces measured at the indicated ROIs (C), and average traces per stimulus (D) measured in the head-fixed mice expressing CRF1.0 (B1-D1, top panels) or CRFmut (B1-D1, bottom panel). The mice were subjected to tail shock (A1-D1) forced running on a treadmill (A2-D2), and two-photon imaging was performed in the motor cortex and prefrontal cortex (PFC). Scale bars, 100 μm. (E and F) Representative traces (E) and summary (F) of the rise and decay t_50_ values of the CRF1.0 signal in response to tail shock and forced running; n = 10-12 trials from 3 mice. (G) Summary of the peak fluorescence response measured in the motor cortex and PFC in mice expressing CRF1.0, CRFmut, or EGFP-CAAX in response to tail shock and forced running; n = 3-7 mice each.

Finally, head-fixed mice were forced to run on a treadmill. In response to this stressful stimulus, CRF1.0 fluorescence was monitored using 2-photon microscopy (Figure 8A2). We found that at the onset of forced running, CRF1.0 fluorescence first increased, then reached a plateau within approximately 5 s, and finally returned to baseline after the treadmill stopped; in contrast, no response was measured in mice expressing CRFmut or EGFP-CAAX (Figure 8B2-D2, G and Figure S6A-E; the kinetics and time constants are shown in Figure 8E-F). These results suggest that the CRF1.0 sensor is sufficiently sensitive to detect the endogenous release of CRF *in vivo*.

## DISCUSSION

Here, we report the development and characterization of a series of highly selective and sensitive genetically encoded neuropeptide sensors. Moreover, as proof-of-principle, we show that our SST, CCK and CRF sensors can be used to monitor their corresponding peptides *in vitro*, *ex vivo*, and *in vivo*. For example, we used our SST sensor to monitor activity-dependent SST release in cultured cortical neurons as well as pancreatic islets. In acute brain slices, our CCK and CRF sensors reliably reported the electrical stimulation‒evoked release of CCK and CRF in the hippocampus and the central amygdala, respectively. Moreover, the CRF sensor was successfully used to measure *in vivo* changes in CRF levels in response to stress-inducing stimuli.

Using our peptide sensors, we observed electrically evoked CCK and CRF release in acute brain slices and measured their average apparent diffusion coefficients during signal propagation. This signal spread may derive both from the increasing of peptide release and from the diffusion of released peptides. So, our calculated diffusion coefficients are relatively higher than those of glutamate in the synaptic cleft (∼330 μm^2^/s) (Nielsen et al., 2004), dopamine in the rat brain (∼68 μm^2^/s) (Rice et al., 1985), and GFP-tagged tissue plasminogen activator (∼0.02 μm^2^/s) (Weiss et al., 2014) estimated by other methods. Further studies could apply optogenetic and chemogenic tools to drive the release from peptidergic neurons. By combining these GRAB peptide sensors with neurotransmitter sensors, we anticipate that it is possible to monitor the real-time release of both neuropeptides and neurotransmitters, providing new insights into the mechanisms and functions of neuropeptide corelease.

In addition to its use *in vitro* in cultured neurons, we also measured the endogenous SST release in isolated pancreatic islets, consisting of cell types that secrete glucagon and insulin to maintain blood glucose levels (Hauge-Evans et al., 2009). The finest temporal resolution of pulsatile SST release measured in previous studies was on the order of 30 s (Hellman et al., 2012; Salehi et al., 2007). Using our SST sensor, we measured changes in SST levels in response to high glucose at the single-cell level with high temporal resolution on the order of seconds. SST released from δ cells functions as a paracrine regulator to integrate signals from ghrelin, dopamine, acetylcholine, and leptin (Adriaenssens et al., 2016; DiGruccio et al., 2016; Lawlor et al., 2017). Moreover, pancreatic islets receive regulatory input that affects Ca^2+^ fluctuations in α and β cells. These fluctuations are subsequently translated into the appropriate release of glucagon and insulin (Huising et al., 2018). Thus, our SST sensor and other hormone and/or transmitter sensors such as ghrelin, UCN3, DA, and ATP sensors can be combined with Ca^2+^ indicators to study pancreatic islets in healthy conditions and in diabetic animal models.

Finally, our *in vivo* experiments show that these sensors can be used to directly monitor neuropeptide release within specific brain regions during behaviors, supporting their utility in freely moving animals. Although the peptide-expressing cortical neurons are well established (Smith et al., 2020), it’s still intriguing when these peptides are released in a behaviorally relevant manner. In addition to the axonal release, neuropeptides can also be released from LDCVs in the somatodendritic compartment, likely contributing to volume transmission and exerting their function via paracrine modulation (Ludwig and Leng, 2006; Persoon et al., 2018; van den Pol, 2012). Although sensor fluorescence doesn’t directly represent endogenous receptor activation, when and where these neuropeptides are released can therefore be examined using these sensors, thus helping elucidate their regulatory role on neural circuits.

Most of peptide sensors show minimal downstream coupling; for example, the SST1.0 and CCK1.0 sensors exhibit virtually no coupling (Figure 3E-F), suggesting the expression of these peptide sensors will not affect the normal functions of cells. However, the CRF1.0 sensor still shows significant cAMP coupling, albeit with orders of magnitude lower affinity and 60% reduced efficacy (Figure 3E3). The structures of peptide GPCRs bound to G proteins and β-arrestin have been solved, and the interaction sites have been identified (Huang et al., 2020; Liang et al., 2020; Liu et al., 2021; Ma et al., 2020; Yin et al., 2019; Zhang et al., 2021); altering these sites in the CRF1.0 sensor will allow future modifications to further reduce downstream coupling.

In summary, this series of new GRAB peptide sensors can be used both *in vitro* and *in vivo* to monitor the rate and range of peptide release with a high spatiotemporal resolution. These tools promise to advance our understanding of the roles of neuropeptides in health and disease.

## ACKNOWLEDGMENTS

This research was supported by the Beijing Municipal Science & Technology Commission (Z181100001318002 and Z181100001518004); grants from the National Natural Science Foundation of China (31925017, 31871087, and 81821092); grants from the NIH BRAIN Initiative (1U01NS113358 and 1U01NS103558), the Shenzhen-Hong Kong Institute of Brain Science (NYKFKT2019013); the Feng Foundation of Biomedical Research; and grants from the Peking-Tsinghua Center for Life Sciences and the State Key Laboratory of Membrane Biology at Peking University School of Life Sciences to Y.L. We thank Yi Rao for sharing the two-photon microscope. We thank Xiaoguang Lei at PKU-CLS, the National Center for Protein Sciences at Peking University and State Key Laboratory of Membrane Biology at Tsinghua University for providing support for the Opera Phenix high-content screening system. We thank members of the Li lab for helpful suggestions and comments on the manuscript.

## AUTHOR CONTRIBUTIONS

Y.L. designed and supervised the project. H.W. performed the experiments related to developing, optimizing, and characterization neuropeptide sensors in cultured cells, with contributions from T.Q., Y.Y., S.F., G.Lan, G.Li and L.W. Y.Zhao and T.Q. performed the two-photon imaging of sensors in acute brain slices. C.W. and Y.Zhuo performed the *in vivo* two-photon imaging of mice cortex. H.R. and L.X. performed the experiments related to pancreatic islets under the supervision of L.C. and C.T. T.O., L.M. and Y.J. performed *in vivo* ICV infusion and fiber photometry recording experiments under the supervision of D.L. P.C. performed *in vivo* fiber photometry recording experiments under the supervision of J.-N.Z. All authors contributed to the interpretation and analysis of the data. H.W. and Y.L. wrote the manuscript with input from all coauthors.

## LEGENDS

**Figure S1.**
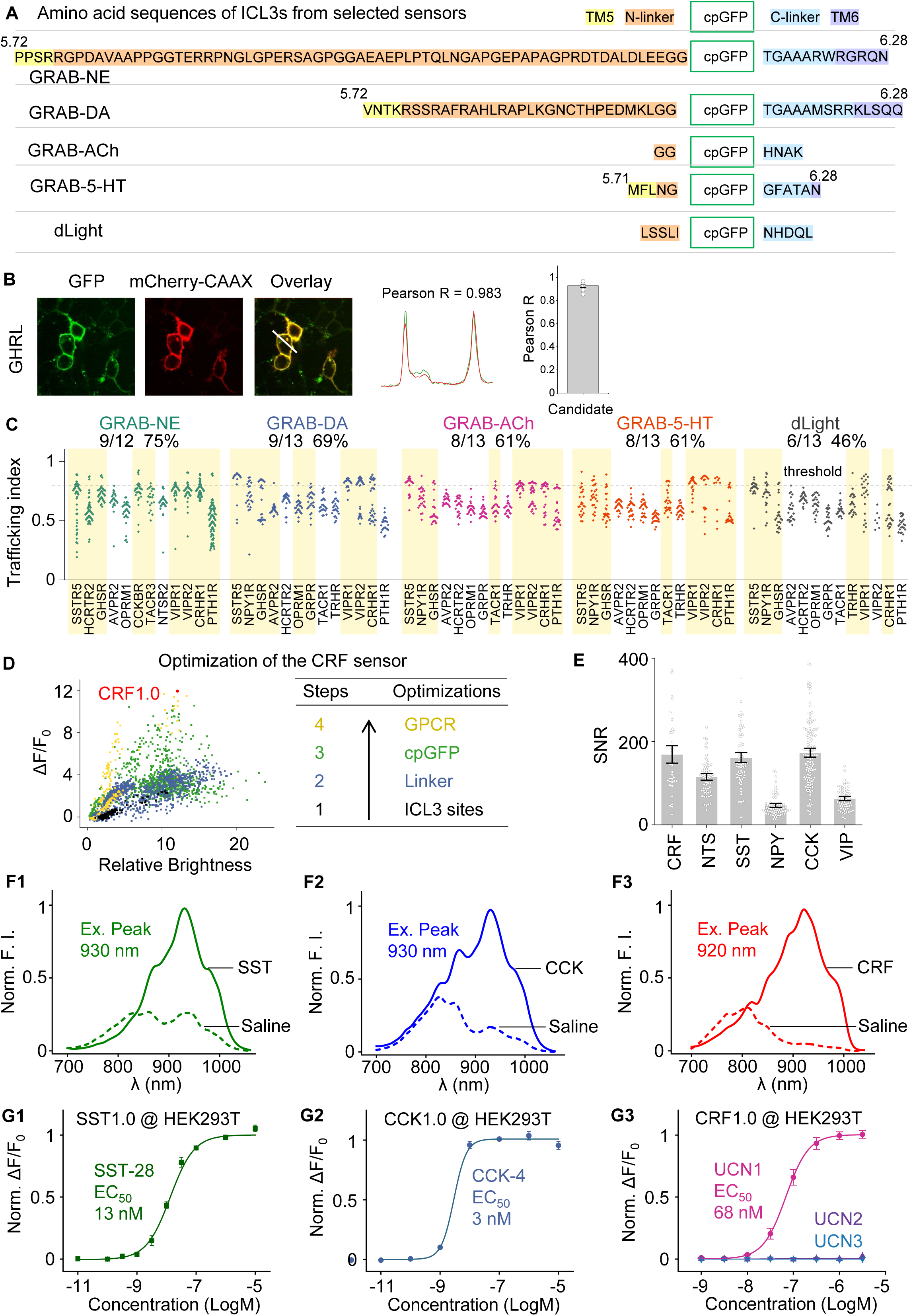
Optimization of GRAB sensors for detecting NTS, SST, NPY, CCK, and VIP. (A) Amino acid sequences of the ICL3 domains in the GRAB-NE, GRAB-DA, GRAB-ACh, GRAB-5-HT, and dLight sensors. (B) Example fluorescence images and intensity line scan profiles of a chimeric GPCR (ghrelin receptor, GHSR) grafted with dLight ICL3 (green), mCherry-CAAX (red), and merged image in the presence of ghrelin (1 μM). The white line indicated the ROI for intensity profiling, and Pearson R was calculated. The averaged Pearson R was used to indicate the membrane trafficking index of sensor variants. (C) Summary of the membrane trafficking index measured for peptide GPCR chimeras containing the ICL3 transplanted from the indicated sensors. The number and percentage of GPCR chimeras with a maximum trafficking index >0.8 (dashed horizontal line) are shown. Each data point represents the average of 100-300 cells measured in one well. (D) ΔF/F_0_ plotted against relative brightness of CRF sensor candidates during the 4-step optimization process. CRF1.0 was the candidate sensor with the highest response and relatively high brightness and is indicated by the red dot. (E) Summary of the signal-to-noise ratio (SNR) of the CRF1.0, NTS1.0, SST1.0, NPY1.0, CCK1.0, and VIP1.0 sensors expressed in HEK293T cells in response to 1 μM of the corresponding ligand; n = 41-119 cells from 3-4 cultures. (F) Two-photon excitation cross-sections of the SST1.0 (F1), CCK1.0 (F2), and CRF1.0 (F3) sensors expressed in HEK293T cells in the presence of saline (dashed lines) or the corresponding ligand (solid lines). Normalized fluorescence intensity is plotted on the *y*-axes. (G) Normalized dose-response curves of the change in SST1.0 (G1), CCK1.0 (G2), and CRF1.0 (G3) fluorescence in response to the indicated concentrations of somatostatin-28 (SST-28), CCK-4, urocortin 1 (UCN1), UCN2, and UCN3; n = 3 wells containing 100-300 cells per well.

**Figure S2.**
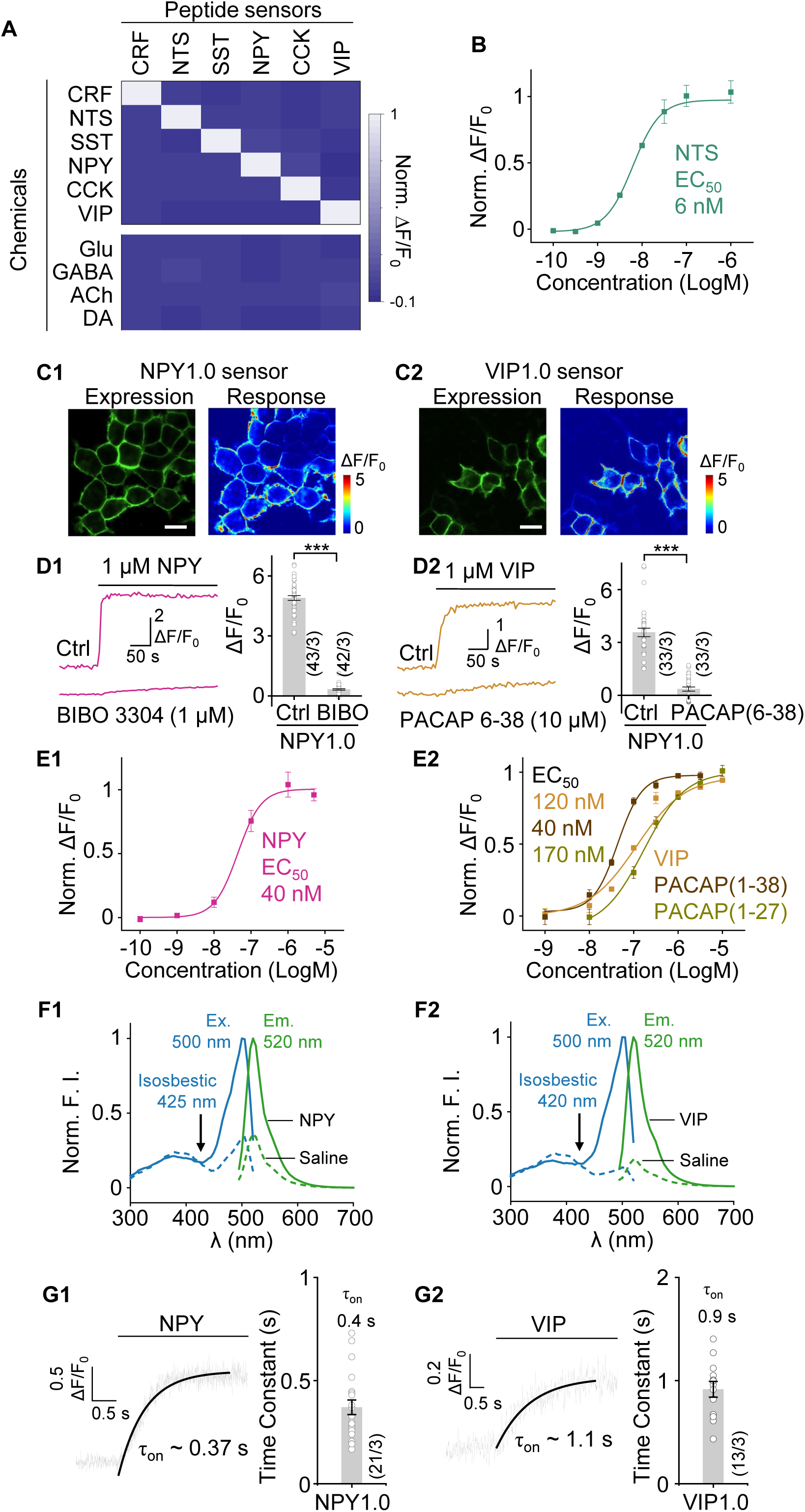
Characterization of NTS1.0, NPY1.0, and VIP1.0 expressed in HEK293T cells. (A) The normalized change in fluorescence for all six peptide sensors were measured in response to the application of the indicated compounds. CRF, neurotensin (NTS), SST-14, neuropeptide Y (NPY), CCK-8s, and vasoactive intestinal peptide (VIP) were applied at 10 μM, while glutamate (Glu), γ-aminobutyric acid (GABA), acetylcholine (ACh), and dopamine (DA) were applied at 10 μM. n = 4 wells containing 100-300 cells per well. (B) Normalized dose-response curve for NTS1.0-expressing HEK293T cells in response to the indicated concentrations of neurotensin; n = 3 wells containing 100-300 cells per well. (C) Representative expression and pseudocolor responses of NPY1.0 (C1) and VIP1.0 (C2) expressed in HEK293T cells in response to the corresponding ligands. Scale bars, 20 μm. (D) Example traces (left) and summary (right) of NPY1.0 (D1) and VIP1.0 (D2) fluorescence measured in HEK293T cells pre-incubated with either saline (Ctrl) or the indicated antagonist; where indicated, NPY and VIP were applied. n = 33-44 cells in 3 wells. (E) Normalized dose-response curves for NPY1.0 (E1) and VIP1.0 (E2) expressed in HEK293T cells in response to the indicated concentrations of the indicated ligands; n = 3 wells containing 100-300 cells per well. (F) Excitation (blue traces) and emission (green traces) spectra of NPY1.0 (F1) and VIP1.0 (F2) expressed in HEK293T cells measured in saline (dashed lines) or ligand (1 μM NPY or 10 μM VIP; solid lines). (G) Representative change in NPY1.0 (G1) and VIP1.0 (G2) fluorescence cells in response to local perfusion of 10 μM NPY or 1 mM VIP, imaged using confocal line scanning. The data were fitted to obtain τ_on_. Each representative trace is the average of 3 ROIs on the scanning line, fitted with a single-exponential function (black curves). Shown at the right is the summary data; n = 13-21 cells from 3 cultures.

**Figure S3.**
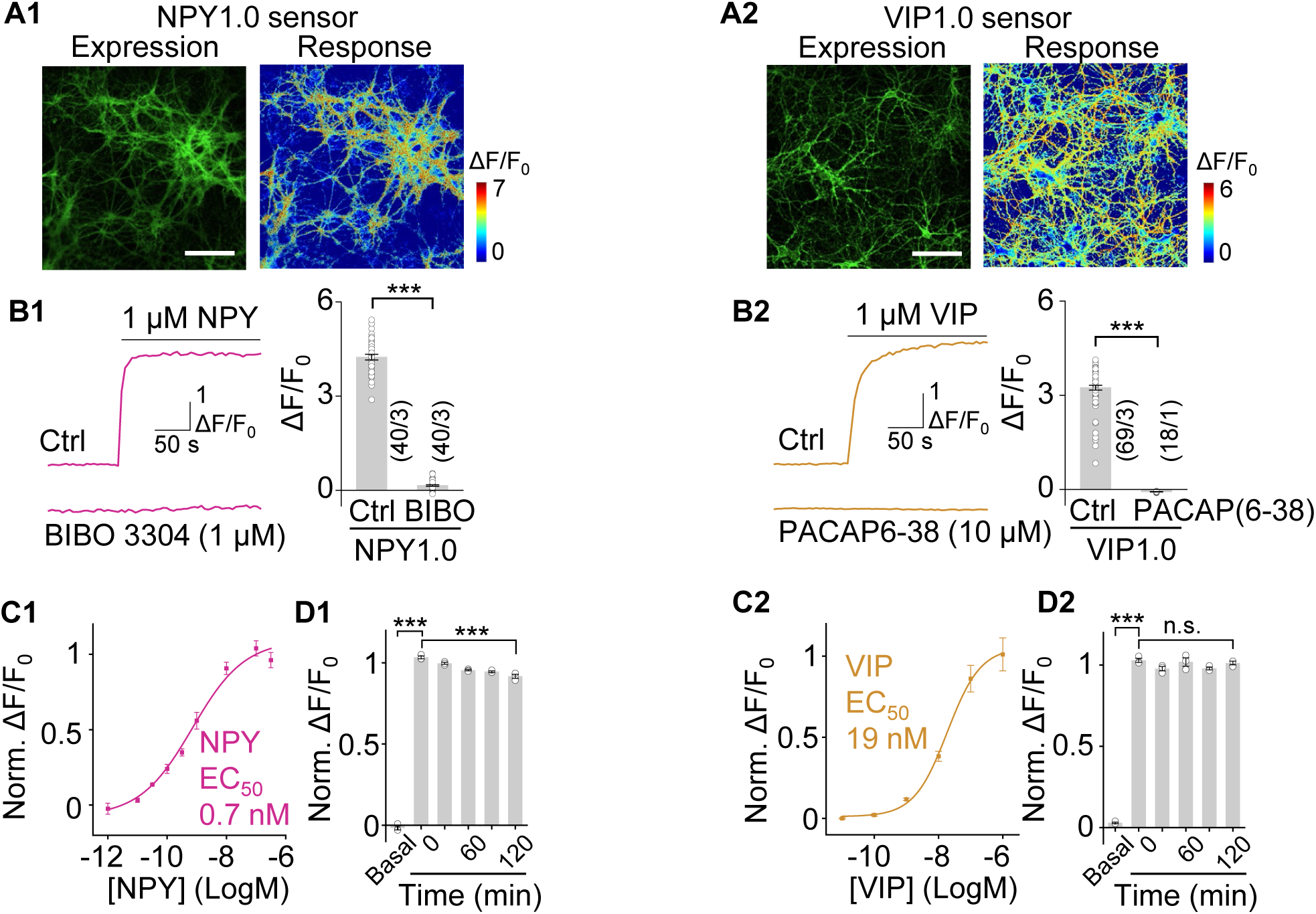
Characterization of NPY1.0 and VIP1.0 expressed in cultured neurons. (A) Expression (left) and pseudocolor responses (right) of primary cultured rat cortical neurons expressing NPY1.0 (A1) or VIP1.0 (A2) in response to 1 μM NPY and 1 μM VIP, respectively. Scale bars, 100 μm. (B) Representative fluorescence traces and summary responses of neurons expressing NPY1.0 (B1) or VIP1.0 (B2) in response to 1 μM NPY or 1 μM VIP, respectively; where indicated, the neurons were pre-incubated with 1 μM BIBO 3304 or 10 μM PACAP (6-38). n = 18-69 ROIs from 1-3 cultures. (C) Normalized dose-dependent fluorescence responses measured in cells expressing NPY1.0 (C1) and VIP1.0 (C2); n = 3 cultures containing 20-40 neurons per culture. (D) Normalized change in NPY1.0 (D1) or VIP1.0 (D2) fluorescence before and after continuous application of 100 nM NPY or 1 μM VIP, respectively, for up to 120 minutes; n = 3-4 cultures containing 20-40 neurons per culture.

**Figure S4.**
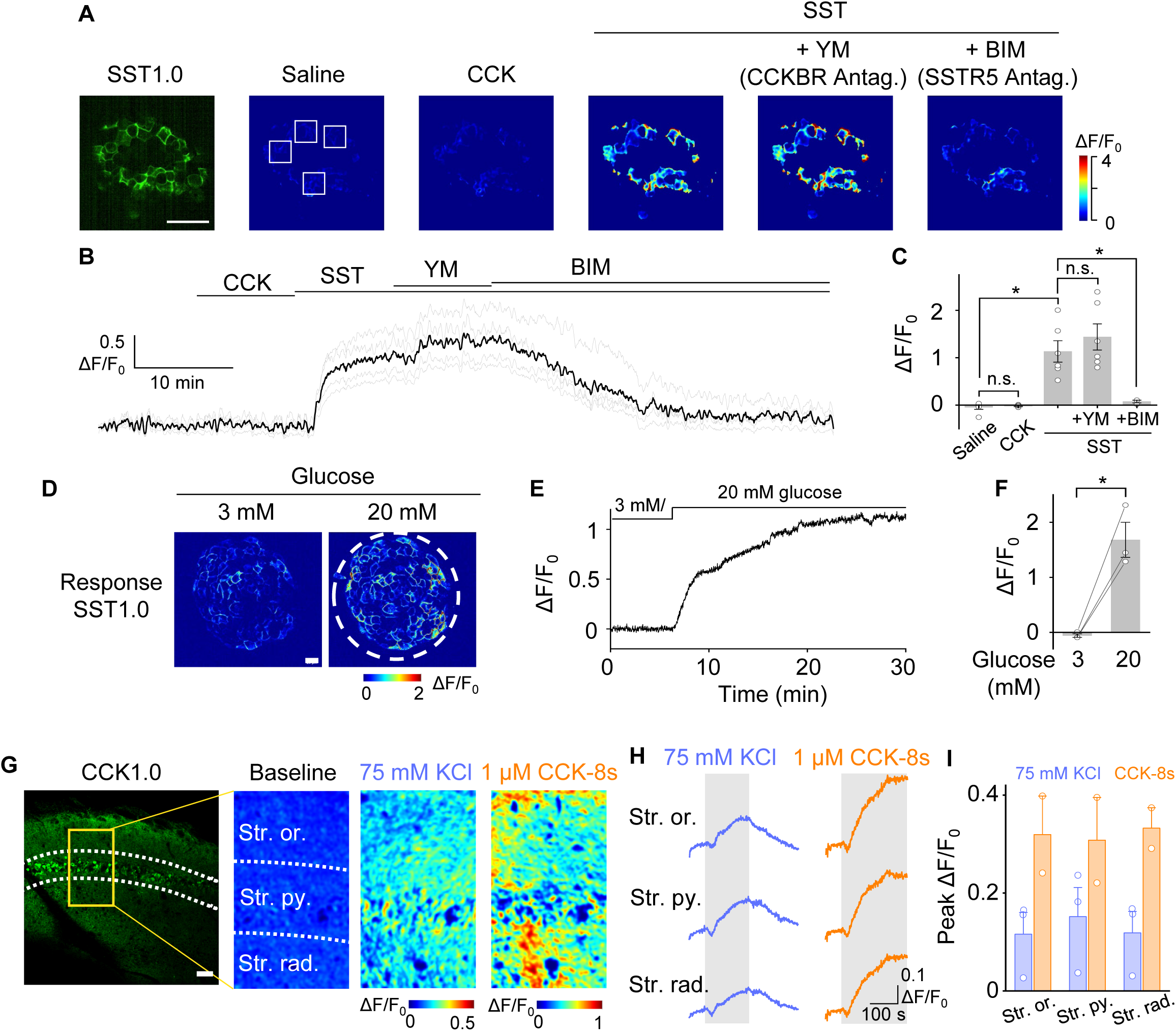
Validation of the SST1.0 sensor in pancreatic islets and validation of the CCK1.0 sensor in acute hippocampal slices. (A) Expression and pseudocolor responses measured in a mouse pancreatic islet expressing SST1.0 in control solution (saline supplemented with 250 μM diazoxide), 1 μM CCK-8s, and 1 μM SST-14 in the absence and presence of the CCKBR antagonist YM022 (10 μM) or the SSTR5 antagonist BIM23056 (10 μM). White squares indicate the ROIs for quantification. Scale bar, 50 μm. (B) Average (black trace) and raw (gray traces) fluorescence responses measured at the 4 ROIs shown in (B). (C) Summary of the change in SST1.0 fluorescence measured in SST1.0- expressing islets under the indicated conditions; n = 3-6 islets each. (D) Pseudocolor images of a mouse islet expressing SST1.0 in the presence of 3 mM and 20 mM glucose. The dashed circle indicates the islet. Scale bar, 20 μm. (E and F) Representative fluorescence trace (E) and summary data of SST1.0-expressing mouse islets in the presence of 3 mM and 20 mM glucose. (G-I) Representative fluorescence and pseudocolor images (G), example traces (H), and summary peak response (I) of CCK1.0-expressing hippocampal slices in response to 75 mM KCl or 1 μM CCK-8s; n= 2-3 slices per group. Scale bar, 50 μm.

**Figure S5.**
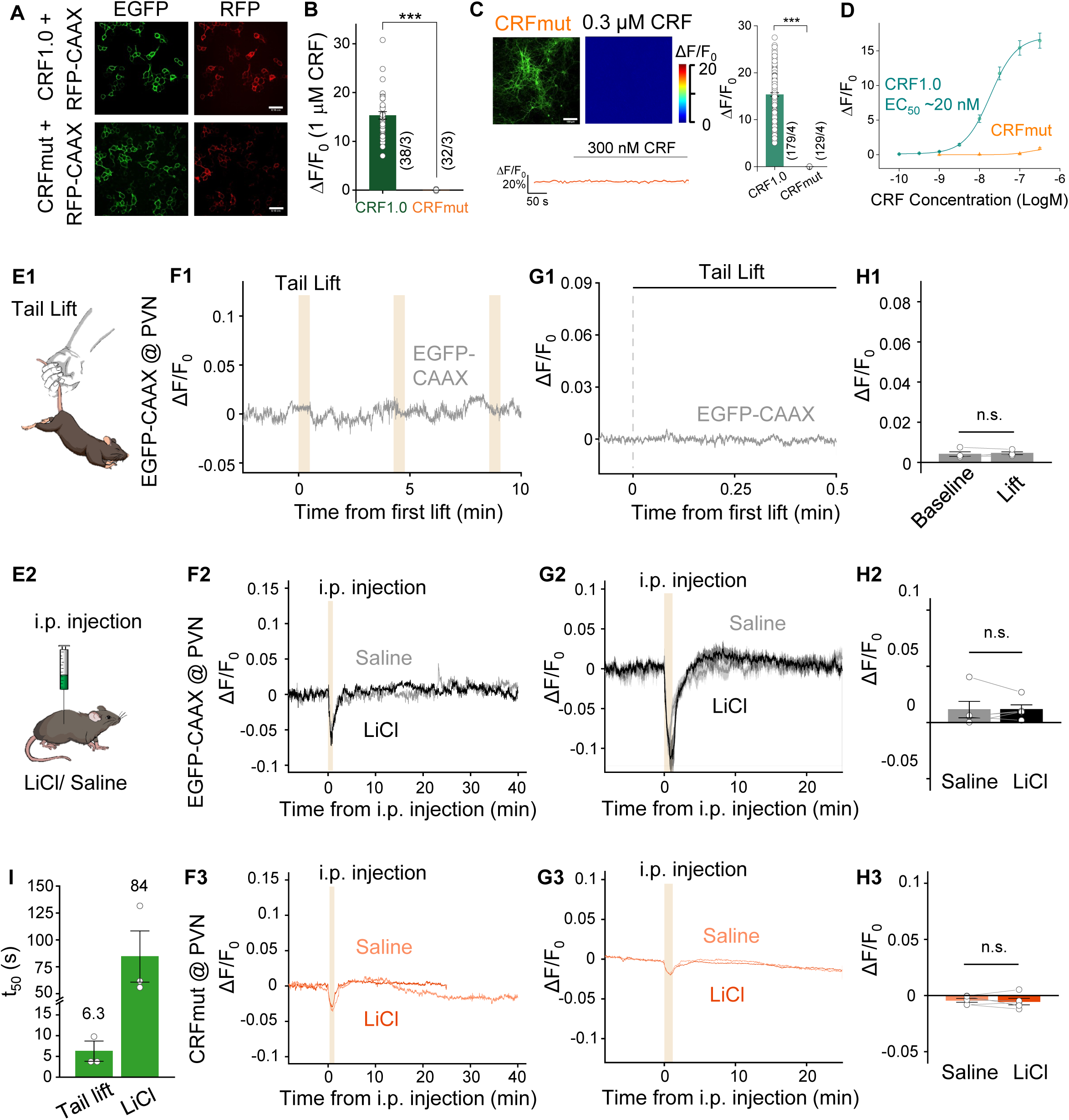
CRF1.0 can detect endogenous CRF release using fiber photometry. (A) Fluorescence images of HEK293T cells co-expressing RFP-CAAX together with CRF1.0 (top row) or CRFmut (bottom row). Scale bars, 100 μm. (B) Summary of the change in fluorescence measured in cells expressing CRF1.0 or CRFmut in response to 1 μM CRF; n = 32-38 cells from 3 cultures. (C) Expression and pseudocolor responses (upper left panel), a representative fluorescence trace (lower left panel), and summary of the response (right panel) measured in rat cortical neurons expressing CRFmut in response to 300 nM CRF; for comparison, the summary data also includes the response measured in neurons expressing CRF1.0 (n = 129-179 ROIs from 4-6 cultures). Scale bars, 100 μm. (D) Dose-response curves measured in neurons expressing CRF1.0 or CRFmut in response to CRF; n = 3 cultures containing 20-40 neurons per culture. (E) Schematic diagram depicting the experimental strategy. CRFmut or EGFP-CAAX was virally expressed in the paraventricular nucleus (PVN); 3 weeks later, the mice received a 30-s tail lift or an i.p. injection of LiCl or saline, and fluorescence was measured in the PVN. (F-H) Representative fluorescence traces (F), average traces (G), and summary data (H) of CRFmut or EGFP-CAAX fluorescence in the PVN following a 30-s tail lift or an i.p. injection of saline or LiCl; n=5 and 4 animals for CRFmut and EGFP-CCAX group, respectively. (I) Summary of the rise t_50_ values of the CRF1.0 signal in response to tail lift and i.p. injection of LiCl; n = 3 mice per group.

**Figure S6.**
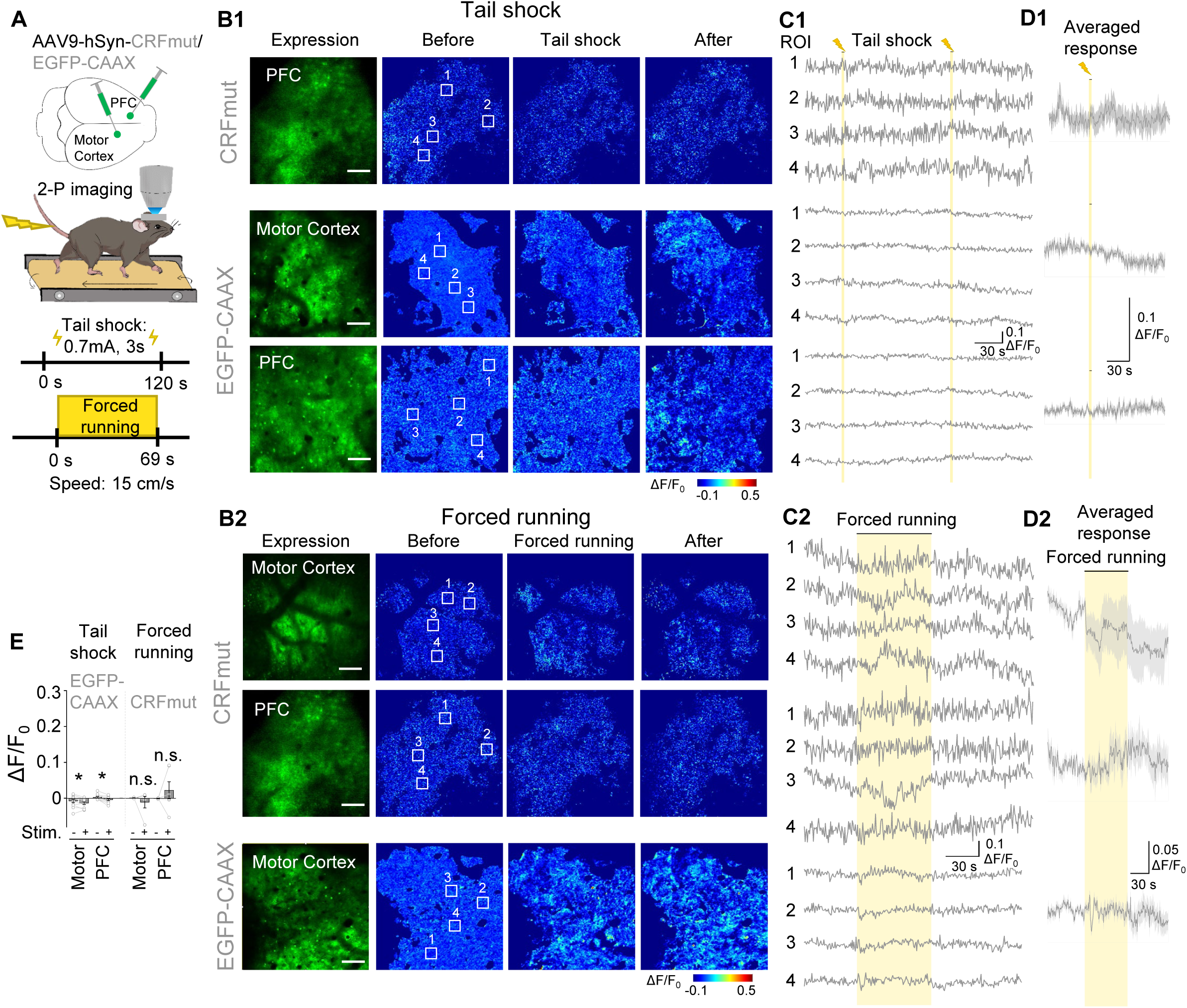
CRF1.0 can detect endogenous CRF release using 2-photon imaging. (A-E). Schematic diagram (A), representative fluorescence and pseudocolor images (B), representative traces of the indicated ROIs (C), average traces (D), and summary responses (E) measured in head-fixed mice expressing either CRFmut or EGFP-CAAX in the motor cortex and prefrontal cortex (PFC). Where indicated, the mice were subjected to the tail shock paradigm (B1-D1) or forced running on a treadmill (B2-D2) during two-photon *in vivo* imaging of the motor cortex and PFC. Scale bars, 100 μm.

**Video S1. Glucose stimulated SST release in isolated pancreatic islets, related to Figure4.**

## METHODS

### Cell lines

HEK293T cells (CRL-3216, ATCC) were used to generate cell lines stably expressing the CRF1.0, SST1.0, CCK1.0, NPY1.0, NTS1.0, and VIP1.0 sensors. These stable cell lines were generated by transfecting cells with pCS7-PiggyBAC (S103P, S509G) (Yusa et al., 2011) together with vectors containing a 5’ PiggyBac inverted terminal repeat sequence (ITR), CAG promoter, the GRAB peptide sensor coding region, IRES sequence, a puromycin-encoding gene, and a 3’ PiggyBac ITR; 24 hours after transfection, the cells were selected by culturing in 1 μg/ml puromycin. The HTLA cell line for the Tango assay was a gift from Bryan L. Roth (Kroeze et al., 2015). All cell lines were cultured in DMEM (Biological Industries, 06-1055-57-1ACS) supplemented with 10% (v/v) fetal bovine serum (FBS; CellMax, SA301.02) and 1% (v/v) penicillin-streptomycin (Gibco, 15140122) at 37°C in humidified air containing 5% CO_2_.

### Cultured rat primary cortical neurons

Rat cortical neurons were obtained from postnatal day 0 (P0) Sprague–Dawley rat pups of both sexes (Beijing Vital River Laboratory Animal Technology Co., Ltd.). In brief, the brain was removed, and the cortex was dissected, dissociated in 0.25% trypsin-EDTA (Gibco, 25200-056), and plated on glass coverslips pre-coated with poly-D-lysine hydrobromide (Sigma, P7280). The neurons were cultured in Neurobasal medium (Gibco, 21103049) supplemented with 2% B-27 (Gibco, A3582801), 1% GlutaMAX (Gibco, 35050061), and 1% penicillin-streptomycin (Gibco, 15140122) at 37°C in humidified air containing 5% CO_2_.

### Mice

C57BL/6N mice (6-8 weeks of age and 10-12 weeks of age) were obtained from Beijing Vital River Laboratory Animal Technology Co., Ltd. and group-housed (up to five mice per cage) under a 12-h/12-h light/dark cycle with the ambient temperature maintained at 25°C. All surgical and experimental protocols were approved by the Animal Care and Use Committee at Peking University, the University of Science and Technology of China, New York University, the Institute of Neuroscience, and the Chinese Academy of Sciences, and were performed in accordance with the standards established by the Association for the Assessment and Accreditation of Laboratory Animal Care.

### Molecular biology

Molecular cloning was conducted using the Gibson assembly method (Gibson et al., 2009). Primers for Gibson assembly were synthesized by Tsingke Biotechnology Co., Ltd., with 30-bp overlap. The coding sequences for the GPCRs were PCR-amplified from the corresponding full-length human GPCR cDNAs (hORFeome database 8.1) using GoldenStar T6 DNA Polymerase (Tsingke, TSE102). The ICL3 from the GRAB-NE (Feng et al., 2019), GRAB-DA (Sun et al., 2020), GRAB-ACh (Jing et al., 2020), GRAB-5-HT (Wan et al., 2021), and dLight (Patriarchi et al., 2018) sensors were PCR-amplified from the corresponding sensors. Chimeric GPCRs and GRAB sensors were cloned into the modified pDisplay vector (Invitrogen) with an upstream IgK sequence and followed by an IRES sequence and mCherry-CAAX. Sanger sequencing was performed to verify the sequence of all clones. GPCR/Sensor-SmBit was constructed from β_2_AR-SmBit, and LgBit-mGs/mGsi/mGsq was a gift from Nevin A. Lambert (Wan et al., 2018). The GRAB peptide sensors were cloned into the pAAV vector under the control of the human *Synapsin* promoter and used for AAV packing.

### Transfection of cell lines and virus infection of primary cultures

HEK293T cells and HTLA cells at 50-60% confluency were transfected with a mixture of polyethylenimine (PEI) and plasmid DNA at a 3:1 (w/w) ratio; after 6-8 h, the transfection reagent was replaced with standard culture medium, and the cells were cultured for an additional 24-36 h for expression of the transfected plasmids.

AAV9 viruses expressing the indicated GRAB peptide sensors were packaged at WZ Biosciences. Each virus (at a titer of 3-5×10^13^ v.g./ml) was added to cultured rat cortical neurons at DIV5-7, and the neurons were imaged 7-10 days later.

### Fluorescence imaging of cultured cells and primary neurons

HEK293T cells and primary neurons were imaged using a Ti-E A1 inverted confocal microscope (Nikon) and an Opera Phenix High-Content Screening System (PerkinElmer). The confocal microscope was equipped with a 10×/0.45 NA objective, a 20×/0.75 NA objective, and a 40×/1.35 NA oil-immersion objective. A 488-nm laser and 525/50-nm emission filter were used to image green fluorescence, and a 561-nm laser and 595/50-nm emission filter were used to image red fluorescence. Cells were cultured on glass coverslips in 24- well plates and imaged in a custom-made chamber. The Opera Phenix system was equipped with 20×/1.0 NA and 40×/1.15 NA water-immersion objectives. A 488-nm laser and 525/50-nm emission filter were used to image green fluorescence, and a 561-nm laser and 600/30-nm emission filter were used to image red fluorescence. Cells were cultured and imaged in CellCarrier Ultra 96- well plates (PerkinElmer).

The cells were imaged in Tyrode’s solutions containing (in mM): 150 NaCl, 4 KCl, 2 MgCl_2_, 2 CaCl_2_, 10 HEPES, and 10 glucose (pH adjusted to 7.35-7.45 with NaOH). Where indicated, the following compounds were applied to the cells in Tyrode’s solution via bath application or a custom-made perfusion system: SST-28 (Anaspec), SST-14 (Anaspec), CCK-8s (Abcam), CCK-4 (Abcam), CRF (Anaspec), UCNI (MedChemExpress), UCNII (MedChemExpress), UCNIII (Abcam), NTS (Anaspec), NPY (Abcam), VIP (Anaspec), PACAP(1-38) (MedChemExpress), PACAP(1-27) (MedChemExpress), Orexin-B (GL Biochem), Substance P (Tocris), Ghrelin (Tocris), Teriparatide (Human parathyroid hormone-(1-34)) (MedChemExpress), Glu (Sigma-Aldrich), GABA (Tocris), DA (Sigma-Aldrich), BIM23056 (Abcam), YM 022 (Tocris), NBI 27914 (Santa Cruz), Antalarmin (Cayman), α-helical CRF (Tocris), SR142948 (Tocris), BIBO 3304 (Tocris), and PACAP(6-38) (Tocris). For high K^+^ stimulation, Tyrode’s solution contained 79 mM NaCl and 75 mM KCl. For screening candidates using SSTR5, NPY1, NPY5, GHSR, AVPR2, NTSR1, CCKBR, HCRTR2 (OX2), TACR1 (NK1), TRHR, VIPR1, VIPR2, CRHR1, or PTH as scaffolds in Figure 1, the following compounds were applied respectively (in μM): 1 SST-28, 1 NPY, 1 NPY, 1 Ghrelin, 5 Desmopressin (Tocris), 1 NTS, 1 CCK-8s, 1 Orexin-B, 1 DAMGO (Tocris), 10 Bombesin (Tocris), 10 Substance P, 20 Taltirelin (Tocris), 1 VIP, 1 VIP, 1 CRF, and 1 Teriparatide.

### Spectra measurements

The linear optical properties of the GRAB peptide sensors expressed in HEK293T cells were measured using a Safire 2 plate reader (Tecan). Cells were harvested and transferred to black-wall 384-well plates containing either saline alone or saline containing the corresponding peptides. Emission spectra were measured using an excitation wavelength of 455 nm with a bandwidth of 20 nm, and emissions were collected using an emission wavelength step size of 5 nm. Excitation spectra were measured using excitation light ranging from 300 nm to 520 nm with a wavelength step size of 5 nm, and emission light was collected at 560 nm with a bandwidth of 20 nm.

The 2-photon fluorescence spectra of the GRAB peptide sensors expressed in HEK293T cells were measured at 10-nm increments from 700-1050 nm using a Bruker Ultima Investigator 2-photon microscope equipped with Spectra-Physics Insight X3. Cells were measured in Tyrode’s solutions or Tyrode’s solutions containing the corresponding peptides. The 2-photon laser power at various wavelengths was calibrated, and the fluorescence measured in un-transfected cells was subtracted as background.

### Tango assay

HTLA cells were cultured and transfected in 6-well plates, placed in 96-well plates (white with a clear flat bottom), and solutions containing various concentrations of peptides were applied; 12 h after induction, the medium was discarded, and 40 μl of Bright-Glo Luciferase Assay Reagent (Promega) diluted 20-fold in phosphate-buffered saline (PBS) was added to each well at room temperature. After a 10-min reaction in the dark, luminescence was measured using a Victor X5 multi-label plate reader (PerkinElmer).

### Mini G protein luciferase complementation assay

HEK293T cells were cultured and transfected in 6-well plates and grown to 80-90% confluency. The cells were then dissociated using a cell scraper, resuspended in PBS, and placed in 96-well plates (white with a clear flat bottom) containing Nano-Glo Luciferase Assay Reagent (Promega) diluted 1000-fold in PBS at room temperature. Solutions containing various concentrations of peptides were added to the wells. After a 10-min reaction in the dark, luminescence was measured using a Victor X5 multi-label plate reader (PerkinElmer).

### Pancreatic islet isolation and imaging of SST1.0 sensor

Male C57BL/6N mice (10 weeks of age) were obtained from Beijing Vital River Laboratory Animal Technology Co., Ltd. The mice were sacrificed by cervical dislocation, and primary pancreatic islets were isolated using collagenase P digestion and purified by hand-picking under a dissecting microscope. After isolation, the islets were cultured overnight in RPMI-1640 medium containing 10% fetal bovine serum (10099141C, Gibco), 8 mM D-glucose, 100 unit/ml penicillin, and 100 μg/ml streptomycin for overnight culture at 37°C in a 5% CO_2_ humidified air atmosphere.

Adenovirus (ADV) expressing the SST1.0 sensor (pAdeno-MCMV-SST1.0) was prepared by OBiO Technology (Shanghai) Corp., Ltd. The islets were infected with pAdeno-MCMV-SST1.0 by 1 h exposure in 200 μl culture medium (approximately 4×10^6^ plaque-forming units (PFU)/islet), followed by addition of regular medium and further culture for 16-20 h before use.

All fluorescence images were acquired using Dragonfly 200 series (Andor) with a Zyla4.2 sCMOS camera (Andor) and the Fusion software. All channels were collected with a 40x/0.85 NA Microscope Objective (Warranty Leica HCX PL APO).

### Fluorescence imaging of peptide sensors in acute brain slices

Male C57BL/6N mice (6-8 weeks of age) were anesthetized with an i.p. injection of tribromoethanol (Avertin; 500 mg/kg body weight), and the AAV9-hSyn-CRF1.0, AAV9-hSyn-CCK1.0, or AAV9-hSyn-EGFP-CAAX virus (300 nl, 3-5×10^13^ v.g./ml, WZ Biosciences) was injected into the left CeA (AP: -1.2 mm relative to Bregma, ML: -2.5 mm relative to Bregma, DV: -4.4 mm from the dura) or the left CA1 (AP: -2.0 mm relative to Bregma, ML: -1.5 mm relative to Bregma, DV: -1.5 mm from the dura) at a rate of 30 nl/min. After 3 weeks to allow for virus expression, the mice were anesthetized with Avertin and perfused with ice-cold oxygenated slicing buffer containing (in mM): 110 choline-Cl, 2.5 KCl, 7 MgCl_2_, 1 NaH_2_PO_4_, 0.5 CaCl_2_, 25 NaHCO_3_, and 25 glucose (pH 7.4). The brains were dissected, and 300-μm thick coronal slices were cut in ice-cold oxygenated slicing buffer using a VT1200 vibratome (Leica). The slices were transferred and allowed to recover for at least 40 min at 34°C in oxygenated artificial cerebrospinal fluid (ACSF) containing (in mM): 125 NaCl, 2.5 KCl, 1.3 MgCl_2_, 1 NaH_2_PO_4_, 2 CaCl_2_, 25 NaHCO_3_, and 25 glucose (pH 7.4). The brain slices were then transferred to a custom-made perfusion chamber and imaged using an FV1000MPE 2-photon microscope (Olympus) or Bruker 2-photon microscope. CRF1.0, CCK1.0, and EGFP-CAAX were excited using a 920-nm 2-photon laser, and electrode tips were placed near the CeA or CA1 region expressing CRF1.0 CCK1.0, or EGFP-CAAX. Electrical stimuli were applied using an S88 stimulator (Grass Instruments), with a stimulation voltage of 5-8 V and pulse duration of 1 ms.

### Two-photon *in vivo* imaging in mice

Female C57BL/6N mice (6-8 weeks of age) were anesthetized with Avertin, and AAV9-hSyn-CRF1.0, AAV9-hSyn-CRFmut, or AAV9-hSyn-EGFP-CAAX (200 nl, full titer, WZ Biosciences) was injected into the motor cortex (AP: 1.0 mm relative to Bregma, ML: 1.5 mm relative to Bregma, DV: -0.5 mm from the dura) and prefrontal cortex (AP: 2.8 mm relative to Bregma, ML: 0.5 mm; DV: -0.5 mm from the dura). A high-speed drill was then used to open a 4 mm x 4 mm square in the skull. After virus injection, craniotomies were installed with a glass coverslip affixed to the skull surface. A stainless-steel head holder was attached to the animal’s skull using dental cement to help restrain the animal’s head and reduce motion-induced artifacts during imaging. The imaging experiments were performed approximately 3 weeks after surgery. An awake mouse with head mounts was habituated for 10 min in the treadmill-adapted imaging apparatus to minimize the stress associated with head restraint and imaging. The motor cortex or prefrontal cortex was imaged 100-200 μm below the pial surface to measure sensor fluorescence.

A Bruker Ultima Investigator 2-photon microscope equipped with Spectra-Physics Insight X3 was used for *in vivo* imaging. A 920-nm laser was used for excitation, and a 490-560-nm filter was used to measure green fluorescence. All experiments were performed using a 16x/0.8 NA objective immersed in saline, and images were acquired at a frame rate of 1.5 Hz. For the forced running model, the running speed was set at approximately 15 cm/s; for the tail shock model, a 0.7-mA shock was delivered for a duration of 3 s.

After imaging, any motion-related artifacts were corrected using the Non-Rigid Motion Correction (*NoRMscorre*) algorithm. The fluorescence time course was measured using ImageJ software by averaging all pixels within the regions of interest (ROIs). ΔF/F_0_ was calculated using the following equation: ΔF/F_0_=[(F-F_0_)/F_0_], in which F_0_ is the baseline fluorescence signal averaged over a 10-s period before the onset of the forced running or tail shock.

### Fiber photometry recording of CRF1.0 with *in vivo* drug application

Male C57BL/6N mice bred at the NYULMC animal facility (10-12 weeks of age) were anesthetized with isoflurane and placed in a stereotaxic frame. AAV expressing hSyn-CRF1.0 or hSyn-CRFmut (Vigene Biosciences) was injected (160 nl per animal) into the PVN (AP: -0.75 mm relative to Bregma; ML: +0.22 mm relative to Bregma; DV: -4.7 mm from the dura). An optical fiber (400-μm diameter) was implanted 150 μm above the virus injection site (either at the time of virus injection or 2 weeks later). At the same time that the optical fiber was implanted, a bilateral cannula (Plastics One) for drug infusion was also implanted in the dorsal 3rd ventricle or the left lateral ventricle. At least 4 weeks after virus injection, fiber photometry recording was performed in the PVN.

Prior to fiber photometry recording, a ferrule sleeve (ADAL1-5, Thorlabs) was used to connect a matching optic fiber to the implanted fiber, and recordings were performed on the head-fixed wheel. For recording, a 390-Hz sinusoidal 488-nm blue LED light (35 mW; M470F1; Thorlabs) driven by a LEDD1B driver (Thorlabs) was bandpass-filtered (passing band: 472 ± 15 nm, Semrock, FF02-472/30-25) and delivered to the brain to excite CRF1.0 or CRFmut. The emission light passed through the same optic fiber, through a bandpass filter (passing band: 534 ± 25 nm, Semrock, FF01-535/50), and into a Femtowatt Silicon Photoreceiver, which recorded the CRF1.0 or CRFmut emission using an RZ5 real-time processor (Tucker-Davis Technologies). The 390-Hz signals from the photoreceiver were extracted in real-time using a custom-written program (Tucker-Davis Technologies) and used to determine the intensity of the CRF1.0 or CRFmut fluorescence signal.

For generating dose-response curves, CRF (C3042, Sigma or AS-24254, Eurogentec) or AHCRF 9-41 (1184, Tocris) was infused into one of the ventricles via the implanted cannula using a syringe (65457-02, Hamilton). For the data shown in Figure 7B-C, 250 nl of CRF diluted to indicated concentrations (4.8, 1.6, 0.5, 0.05 mg/ml) or 250 nl saline was infused; for the data shown in Figure 5D, 100 nl of 1.6 mg/ml CRF and/or 300 nl of 0.25 mg/ml AHCRF 9-41 was infused. CRF was diluted in distilled water, and AHCRF 9-41 was diluted in distilled water containing 0.1 M NH_4_OH.

For the data shown in Figures 7B and 7D, Friedman’s test was performed, followed by Horm correction. For the data shown in Figure 7C, the two-sided paired Wilcoxon signed rank test was performed. The peak values obtained after applying 1.6 mg/ml CRF in Figure 5B and 5D were the average of all trials from each animal.

### Fiber photometry recording of CRF1.0 during behavioral testing

Male C57BL/6N mice (10-12 weeks of age, from River Vital Laboratory) were anesthetized with an i.p. injection of sodium pentobarbital (80 mg/kg body weight) and AAV9-hSyn-CRF1.0 or AAV9-hSyn-EGFP-CAAX (300 nl, 3-5×10^13^ v.g./ml, WZ Biosciences) was injected into the PVN (AP: -0.80 mm relative to Bregma, ML: -0.25 mm relative to Bregma, DV: -4.60 mm from the dura) at a rate of 40 nl/min. The optic fiber (200 μm inner core diameter, 0.37 fiber numerical aperture, Thinkerbiotech) was implanted 0.20 mm above the injection site and sealed with dental cement. After 4-5 weeks (to allow the mice to recover and to allow for virus expression), a Multi-Channel Fiber Photometry Device (Inper, OPT-FPS-410/470/561) was used for recording. Signals were acquired at a frame rate of 50 Hz, with an exposure time of 9 ms, with gain 0, using 470-nm light 470 at 30-40% power.

For the tail lift experiments, the mouse was suspended by the tail 50 cm above the floor for 30 s per trial. Three 30-s tail lift trials were performed at an interval of ∼220 s; the signal recorded 150 s before the first lift was used as the baseline, and the average of the three responses recorded during the 30-s lifts was used as the lift signal.

For LiCl or saline injection, the signal recorded 500 s before injection was recorded as the baseline. The mice were then briefly anesthetized with isoflurane and given an i.p. injection of saline (0.1 ml/10 g body weight) or LiCl (125 mg/kg body weight) dissolved in saline. The signals were recorded for 2400 s after i.p. injection, and the average response measured during the first 1500 s was used as the LiCl/saline signal.

### Quantification and statistical analysis

Summary data with error bars are presented as the mean ± SEM. Except where indicated otherwise, groups were compared using Student’s *t*-test or a one-way ANOVA with post hoc test, and differences were considered significant at *p*≤0.05. Where applicable, \**p*≤0.05, \*\**p*≤0.01, \*\*\**p*≤0.001, and n.s., not significant.

